# Aging induces aberrant state transition kinetics in murine muscle stem cells

**DOI:** 10.1101/739185

**Authors:** Jacob C. Kimmel, Ara B. Hwang, Wallace F. Marshall, Andrew S. Brack

**Affiliations:** Dept. of Biochemistry & Biophysics University of California, San Francisco; Dept. of Orthopedic Surgery University of California, San Francisco

**Author notes:** Computing, Calico Life Sciences, LLC, South San Francisco, CA, 94080. Memphis Meats, Inc. 2930 Domingo Avenue, Berkeley, CA 94705.

**Keywords:** aging, muscle stem cell, single cell RNA-seq, timelapse imaging, state transition, stem cell activation

## Abstract

Murine muscle stem cells (MuSCs) experience a transition from quiescence to activation that is required for regeneration, but it remains unclear if the transition states and rates of activation are uniform across cells, or how features of this process may change with age. Here, we use timelapse imaging and single cell RNA-seq to measure activation trajectories and rates in young and aged MuSCs. We find that the activation trajectory is conserved in aged cells, and develop effective machine learning classifiers for cell age. Using cell behavior analysis and RNA velocity, we find that activation kinetics are delayed in aged MuSCs, suggesting that changes in stem cell dynamics may contribute to impaired stem cell function with age. Intriguingly, we also find that stem cell activation appears to be a random walk like process, with frequent reversals, rather than a continuous, linear progression. These results support a view of the aged stem cell phenotype as a combination of differences in the location of stable cell states and differences in transition rates between them.

**Summary Statement:** We find that aged muscle stem cells display delayed activation dynamics, but retain a youthful activation trajectory, suggesting that changes to cell state dynamics may contribute to aging pathology.

## 1 Introduction

Stem cells play a keystone role in tissue homeostasis and regeneration across multiple mammalian tissues. During normal homeostasis, stem cells in multiple systems maintain a non-cycling, quiescent state [20]. In the event of injury, quiescent stem cells undergo a dynamic process of activation, generating biomass, restructuring cellular geometry, altering cell metabolism [43], and entering the cell cycle to produce progenitor daughters [8]. Impairment of stem cell activation by alteration of external signals or intrinsic stem cell potential is demonstrated to impair regeneration across multiple tissue systems [60, 36]. Likewise, “priming” of activation by systemic signaling factors has been reported to improve regeneration [40].

In muscle stem cells (MuSCs), the activation process is canonically characterized by expression of *Myod1* [23, 56], loss of *Spry1* and *Pax7*, and entry into the cell cycle [47]. Activation kinetics based on these canonical markers have been characterized at the ensemble level using both transcriptional and protein level analysis [56, 13, 19, 26, 59]. However, these population level assays are unable to address some fundamental questions.

Do cells activate linearly with time, or are some portions of the process faster than others? Linear activation dynamics may suggest that the process involves a cumulative component, whereas non-linear dynamics may suggest a switch-like mechanism. How many intermediary states exist in the activation process? A simple two state system may similarly suggest a switch-like activation mechanism, whereas stable intermediary states are suggestive of a multi-stage process. Intermediary activation states such as “*G*_alert_” [40, 41] have been suggested, but the stability of such states remains unclear. In order to answer these questions, we require measurements of activation dynamics in individual cells.

As MuSCs age, the proportion of cells in regenerative states declines, and the overall regenerative capacity of the stem cell pool is greatly diminished, limiting their expansion and self-renewal potential [4, 7]. Age-related decline in regenerative potential has been attributed to differences in cell signaling [15, 3, 12, 6] or proliferative history [11]. These differences in regenerative potential between stem cells are traditionally viewed as the result of differences in the characteristics of stable cell phenotypes [2, 11, 4].

However, aged MuSCs have been reported to show impaired activation in multiple studies, suggesting that a defect in the activation process may also contribute to impaired regeneration [21, 15, 6, 18]. Cast in the language of dynamical systems, differences in regenerative potential could be the result of cells taking different trajectories, or paths, through state space, or the result of cells moving along the same trajectory at different rates. Impaired activation with age may therefore be explained by one of two models, or a combination of the two.

In the first model (Different Paths), the location of cell states is shifted by age, such that aged cells at a particular point in the activation process exhibit different phenotypes than young cells at the same point in the process. This model could be stated as young and aged cells each exhibiting a unique trajectory through state space. In the second model (Different Rates), aged and young cells are largely similar at comparable points in the activation process, but take different amounts of time to reach a given point. In this model, differences in young and aged phenotypes are primarily the result of changes in activation dynamics. This second model could be stated as young and aged cells obeying different laws of motion along a common constrained trajectory in state space (Fig. 1B).

**Figure 1:**
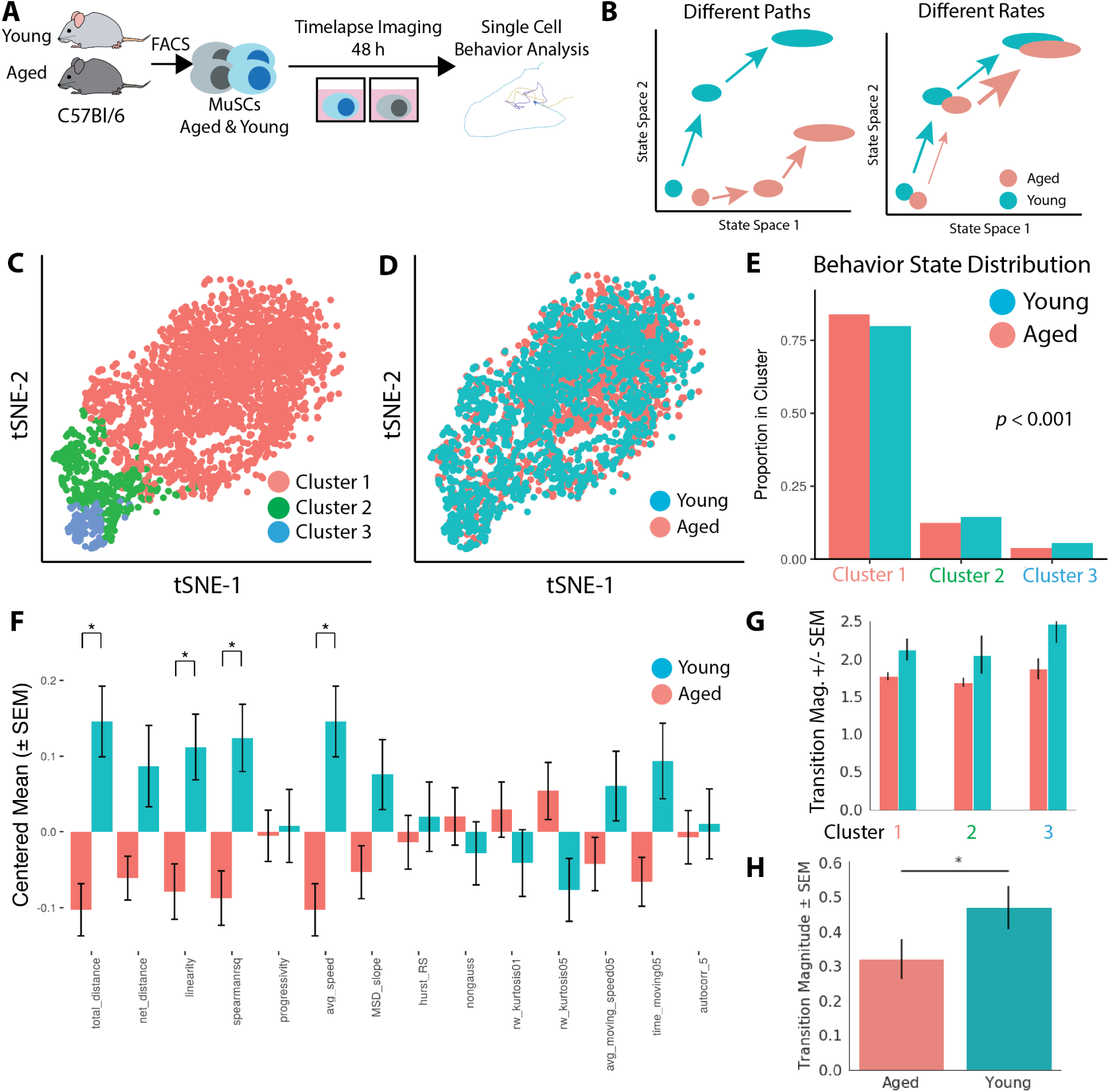
Aged MuSCs display lower cell motility and delayed activation by single cell behavior analysis. **(A)** Experimental schematic. **(B)** Diagram of the Different Paths and Different Rates models for age-related decline in muscle stem cell regenerative capacity. **(C)** t-SNE visualization of cell behavior state space with color overlay of hierarchical clustering identities (*n* = 742 aged and *n* = 1, 201 young cells.). **(D)** t-SNE visualization of aged and young cell identity in cell behavior space. **(E)** Aged cells display a significant preference for less motile cell behavior states. **(F)** Young cells are significantly more motile than aged cells, suggesting aged cells are delayed in activation. Mean feature values are presented for each age after centering the population mean to *µ* = 0 and scaling the variance to *σ*^2^ = 1. **(G)** Aged cells have lower state transition magnitudes within each behavioral state, suggesting each state is dynamically impaired. **(H)** Aged cells have significantly decreased behavior state transition magnitude when considered in aggregate (*t*-test, *p <* 0.05), suggesting delayed activation. State transition magnitude in behavior space is measured as the mean transition vectors magnitude among a group of cells.

Single cell analyses in the hematopoietic system identifies distinct aged and young transcriptional phenotypes (as in the Different Paths model) and altered cell cycle kinetics (as in the Different Rates model) [30], suggesting both models are plausible in the context of myogenic activation. Do aged MuSCs exhibit distinct phenotypes at each stage of activation? Are differences in state transition dynamics in part responsible for diminished regenerative phenotypes in aged MuSCs?

Each of the above questions calls for measuring rates of phenotypic change in MuSCs in addition to measuring cell phenotypes at static timepoints. To measure rates of change, we leverage our recently developed cell behavior analysis platform “Heteromotility” [28] to quantify phenotypic state dynamics during MuSC activation in aged and young MuSCs. Multiple groups have recently demonstrated the value of single cell RNA sequencing (scRNA-seq) to elucidate differences between skeletal muscle cell types and dynamic regulation of myogenic programs following injury [46, 22, 16]. We likewise complement our behavioral assay approach with scRNA-seq sequencing to map the transcriptional state space of MuSC activation. Leveraging RNA velocity analysis [31], we infer transcriptional state transition dynamics to pair with state transition dynamics inferred from cell behavior.

In these transcriptional assays, we further investigate differences across age and activation state within the subsets of highly regenerative label retaining cells (LRCs) and less regenerative non-label retaining cells (nonLRCs). We previously described LRCs and nonLRCs as discrete populations of MuSCs with different proliferative histories and different regenerative potentials [11, 10]. The relative proportions of these populations changes with age, suggesting that age-related changes specific to the LRC or nonLRC compartment may shed light on MuSC aging.

We find that both behavioral and transcriptional state spaces are continuous across MuSC activation and that measurements of cell heterogeneity are comparable between assays. In aged MuSCs, we find aberrant transition dynamics that lead to significantly delayed activation by both methods. These findings are reflected in a comparison of LRCs to less regenerative nonLRCs, suggesting aberrant transition dynamics may be related more generally to impaired regenerative potential. Identifying genetic pathways that are altered in both aged MuSCs and nonLRCs, we find that biosynthetic processes activate more slowly in both populations. To determine if less regenerative MuSCs occupy different steady states, we trained machine learning classifiers to discriminate MuSC age and LRC status. Classifiers readily discriminate between MuSC ages and proliferative histories. Our results suggest that aged stem cells display delayed activation kinetics, in addition to subtle differences in the position of activation states.

## 2 Results

### 2.1 Activation kinetics are delayed in aged MuSCs

Previously, we have demonstrated that quantitative measurements of cell motility behavior from timelapse imaging data are sufficient to resolve states of MuSC activation and state transitions [28]. This approach allows for the direct observation of cell state transitions during MuSC activation and quantitative measurement of transition rates. Cell behavior measurements also have inherent functional relevance in the MuSC context, where motility behaviors are necessary for cells to translocate to the site of injury signals.

We applied this cell behavior analysis method to aged and young MuSCs to determine (1) if aged cells occupied distinct behavioral states and (2) if aged cells exhibit different cell state transition dynamics during activation. MuSCs were isolated from aged (20 m.o., *n* = 1) and young (3 m.o. *n* = 1) mice by FACS and cultured sparsely in 96-well plates with rich growth media to stimulate MuSC activation (see Methods). Timelapse imaging was performed using an automated, incubated microscopy platform for 48 hours after plating, with images taken every 15 minutes (Fig. 1A). This temporal window captures the early stages of MuSC activation, including the switch from a quiescent *Pax7+*/*Myod1-* state to a *Myod1+* state [13].

We quantified cell behaviors across the latter 33 hours of the timelapse using Heteromotility (see Methods). Visualizing cell behavior state space with t-SNE [52] reveals heterogeneous cell behavior states, previously shown to reflect different states of MuSC activation (Fig. 1C) [28]. Performing unsupervised hierarchical clustering to identify cell behavior states reveals that three cell behavior states optimizes the Silhouette index (color labels in Fig. 1C, see Methods). Cluster 1 is largely immotile, Cluster 3 displays limited motility behavior, and Cluster 2 displays more extensive and dramatic motility behaviors.

Aged and young cells do not occupy distinct regions in behavioral state space (Fig. 1D). This suggests that aged and young cells share a common set of behavioral states, but aging may induce preferences in state occupancy rates. Quantifying the proportion of aged and young cells in each motility state revealed that aged cells preferentially occupy the less motile behavior states relative to young cells (Fig. 1E). This preference for less motile states was significant, as assessed by the *χ*^2^ test of the Age × Behavior State contingency table (*p <* 0.001). As motility is associated with activation, a preference for less motile states among aged cells suggests they exhibit a slower phenotypic change from quiescence to activation.

Comparing individual cell behavior features between aged and young cells confirms that aged cells are significantly less motile than young cells. Metrics of total motility distance, motility linearity, and average motility speed are all significantly higher in young cells. Metrics of kurtosis and autocorrelation are also significantly increased in young cells (*t*-test, *p <* 0.05, Holm-Bonferroni corrected) (Fig. 1F, note that feature values are zero-centered and scaled to unit variance).

Do aged cells transition differently between motile states, in addition to having a preference among them? To answer this question directly, the state transition rates across aged and young cells were quantified. Quantification was performed by splitting the time course into 4 adjacent windows, each *τ* = 5 hours in length. Cell state was defined within each *τ* length window as a location in 2D PCA space. State transitions were measured for each cell as the distance between each sequential pair of cell states (see Methods). Aged cells had lower state transition magnitudes (Fig. 1G, H, *p <* 0.05 *t*-test).

This result indicates that aged cells are delayed in activation relative to young counterparts. This is reflected by the enrichment of aged cells in less activated behavioral states and dampened state transition rates we observe in aged cells. We interpret this result as support for the Different Rates model of MuSC aging noted above, where aberrant dynamics between states contribute to the functional defects observed in aged MuSCs.

### 2.2 Transcriptome analysis reveals progressive states of MuSC activation

Cell behavior analysis allows for inference of cell state at the phenotypic level, but does not provide direct insight into the molecular determinants of different cell states. To generate a portrait of the molecular state space of activating MuSCs, we employed single cell RNA-sequencing (scRNA-seq). In addition to determining the transcriptional differences underlying different states of MuSC activation, we sought to determine how the dynamics of activation differ across MuSC populations with different regenerative capacity.

To this end, we transcriptionally profiled MuSCs isolated from both young (3 m.o., *n* = 2) and aged (20 m.o., *n* = 2) *H2B-GFP^+/-^;rtTA^+/-^* mice. We previously showed that label-retaining cells (LRCs), which have done fewer divisions during development, are more regenerative than non-label retaining cells (nonLRCs) that have done many divisions [10, 11] The *H2B-GFP^+/-^;rtTA^+/-^* alleles allowed us to isolate MuSCs from these distinct proliferative histories in both young and aged mice by FACS. To capture different states of MuSC activation within these populations, we profiled transcriptomes at two time points. (1) Freshly-isolated “quiescent” MuSCs were processed for scRNA-seq immediately after FACS isolation, and (2) activated MuSCs were processed for scRNA-seq after 18 hours in culture (Fig. 2A). Recent studies have reported that FACS isolated cells used here experience some markers of early MuSC activation, even immediately after isolation. Therefore, it should be noted that our “quiescent,” cell populations experience some early activation stimulus [53, 34]. Our experiment therefore examines three factors: cell age, proliferative history, and activation state (time in culture, 0 hr or 18 hr).

**Figure 2:**
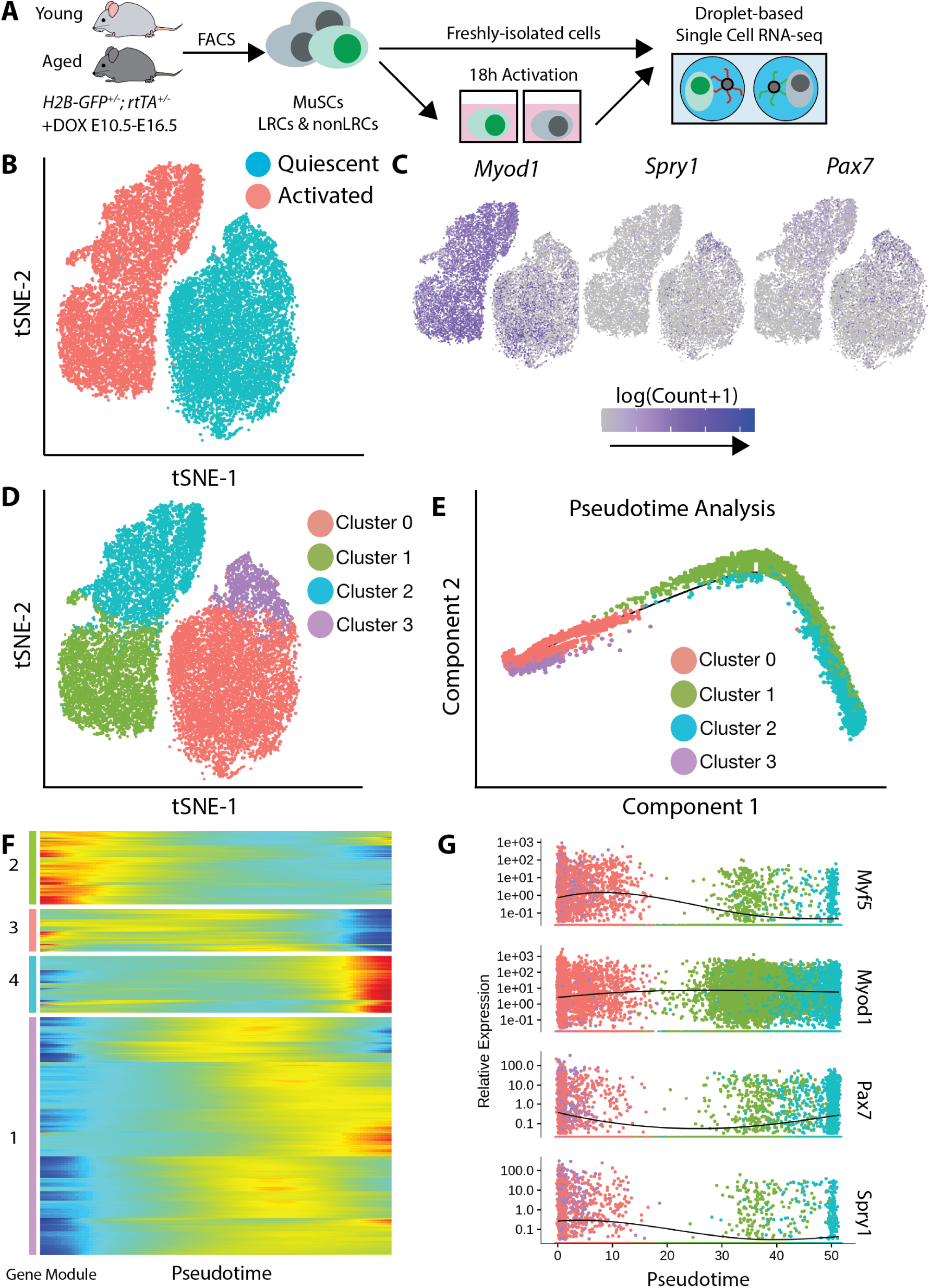
Single-cell RNA-sequencing reveals heterogeneous transcriptional states during myogenic activation. **(A)** Experimental schematic. **(B)** t-SNE visualization of quiescent and activated cells. **(C)** Overlay of MRFs on t-SNE plots to show activated MyoD+ cells localize in a terminal state. **(D)** Definition of heterogeneous transcriptional states by unsupervised clustering. **(E)** Pseudotime analysis of MuSC activation, correctly recapitulating the sequence of ground truth time points. **(F)** Hiearchical clustering identifies four patterns of pseudotemporal gene expression during MuSC activation. **(G)** Visualizing MRF levels across pseudotime reveals that *Pax7* does not decrease monotonically.

After library preparation, sequencing, and quality control, 21,555 individual MuSC transcriptomes were captured (see Methods). This pool captured 8,312 LRCs, 13,243 nonLRCs, 13,927 young cells, 7,628 aged cells, 10,826 quiescent cells, and 10,729 activated cells. A table of cell counts for each condition is provided (Table S1).

We find two discrete clusters of cells in transcriptional space. These discrete clusters correspond to the freshly-isolated and 18h activated cell time points, which we refer to as quiescent and activated cells (Fig. 2B). Each of these discrete clusters contains aged and young LRCs and nonLRCs in a continuous subspace, suggesting that myogenic activation state is a more prominent transcriptional phenotype than either MuSC aging or proliferative history (Fig. S1). Quantifying the proportion of variance explained by each factor using linear models confirms this qualitative observation. Activation accounts for ≈ 12% of variation across transcriptomes, while aging and proliferative history (LRC status) account for *<* 2% (Fig. S1, see Methods).

The expression of characteristic myogenic genes within the transcriptional state space corresponds to known MuSC biology, with the quiescence marker *Pax7* localizing largely to the quiescent cell population and activation marker *Myod1* showing increased expression in the activated cell population (Fig. 2C). However, single cell analysis of these markers on our large sample of MuSCs reveals heterogeneity within both the quiescent and activated populations. Within the quiescent cell population, the subset of *Pax7+* cells occupy the opposite end of quiescent state space from a set of *Myod1+* cells. Likewise, a subset of cells in the activated population express the quiescence marker *Pax7+*.

Given this heterogeneity, we utilized unsupervised Louvain community detection to identify subpopulations within quiescent and activated cells [5]. We tuned the resolution parameter for Louvain clustering to optimize the Silhouette index and find that the optimal resolution (0.2) yields 4 transcriptional clusters. Two clusters lie within the quiescent and activated cell populations respectively (Fig. 2D).

### 2.3 Non-monotonic gene expression patterns are present in myogenic activation

To determine how these transcriptional clusters are temporally ordered during activation, we utilized pseudotime analysis to identify a common pseudotemporal axis through the clusters [38] (see Methods). Pseudotiming infers a distinct sequential ordering for these transcriptional clusters (Fig. 2E). Myogenic gene levels within each of the clusters corroborate the pseudotiming inference with known myogenic biology (Fig. S1). Cluster 3 exhibits the highest levels of quiescence marker *Spry1* and is appropriately selected as the root of the pseudotime axis. Likewise, Clusters 1 and 2 express lower levels of quiescence markers *Pax7*, *Spry1*, and *Cd34*, while expressing higher levels of activation marker *Myod1*.

Pseudotime analysis places Cluster 2 as the end-point of the progression, despite the fact that it contains a subpopulation of *Pax7+* cells while Cluster 1 does not (Fig. S1). This challenges the traditional dogma that *Pax7* levels decrease monotonically with MuSC activation and suggests a more complex temporal regulation of *Pax7* with activation. Fitting splines to *Pax7* expression over pseudotime makes this non-monotonic relationship readily apparent (Fig. 2G). By contrast, the quiescence marker *Spry1* [47] displays a monotonic decrease with activation and *Myod1* displays a monotonic increase (note that the *Myod1* change appears small due to high variance in the population). Previous functional studies with *Pax7* overexpression constructs in MuSCs report that *Pax7* promotes proliferation in certain contexts [58], consistent with increased *Pax7* as cells enter into cycle later in the activation process.

As an orthogonal method to confirm cluster ordering and establish a link between the transcriptional clusters and behavioral clusters, we perform immunostaining following single cell behavior measurements. This analysis finds that the most motile, most activated cell behavior states are enriched for Pax7 protein. This analysis also indicates a non-monotonic regulation for Pax7 across cell behavior states, as we might infer from the ordering of transcriptional clusters (Fig. 3). This result also demonstrates that cell behavior clusters directly reflect molecular features of muscle stem cell activation. More broadly, the similarities we find through these orthogonal assays indicate that cell behaviors contain a high degree of mutual information with transcriptional states and support the use of cell behavior analysis as an orthogonal method to investigate the heterogeneity and dynamism of cell populations.

**Figure 3:**
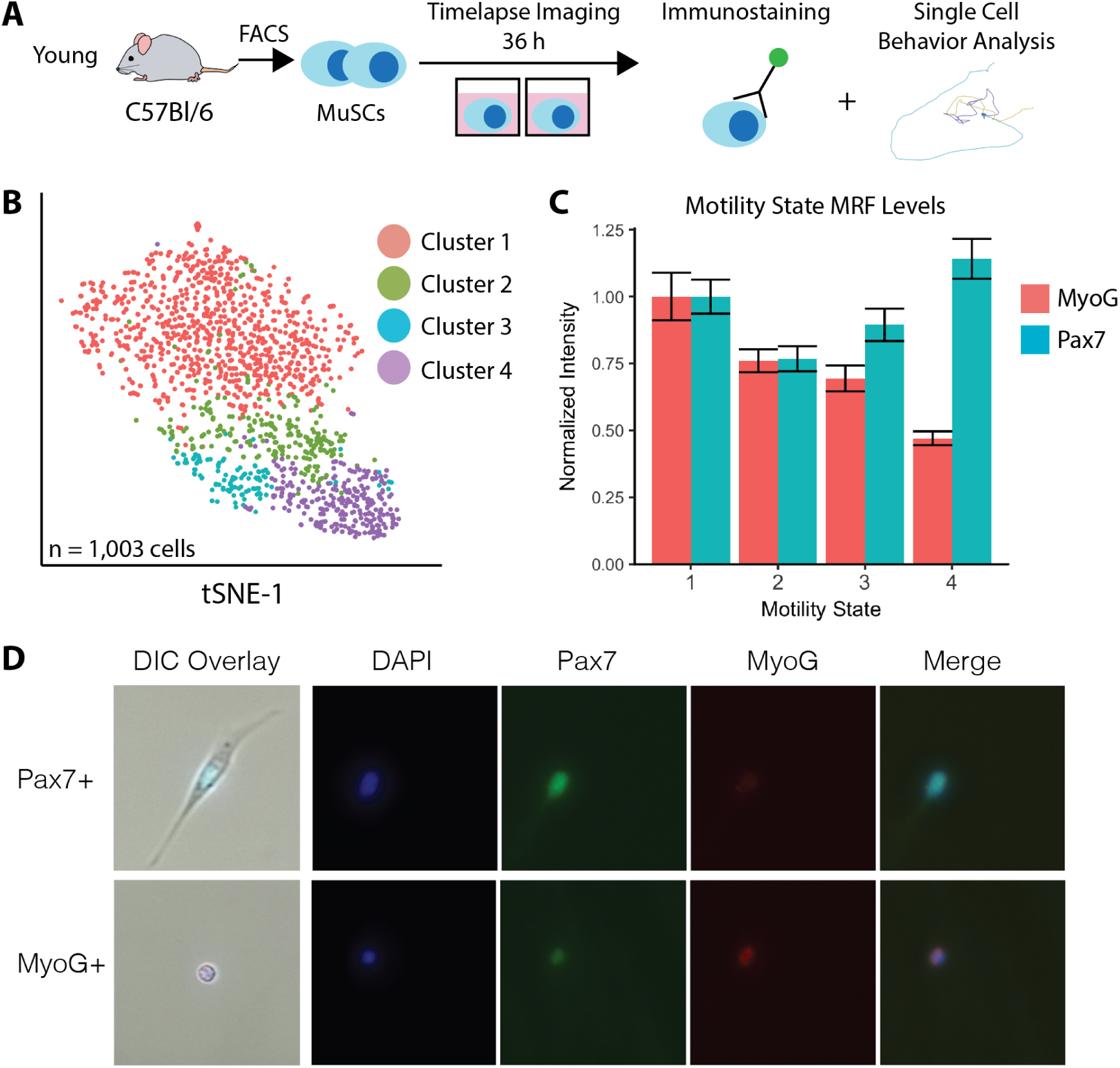
Pax7 is non-monotonically regulated across MuSC cell behavior states during activation. **(A)** Experimental design schematic. MuSCs were isolated, timelapse imaged in culture for 36 hours, and subsequently immunostained. Behavior traces and immunostaining results were matched for each cell by image registration. **(B)** t-SNE visualization of cell behavior states in motility state space, as defined by hierarchical clustering. Behavior state space was generated analyzing 12 hours of tracking data, from 24 hours after isolation to 36 hours. *n* = 1, 003 cells. **(C)** Quantification of immunostaining intensity for Pax7 and MyoG within each cell behavior cluster. Pax7 displays a non-monontic relationship with cell behavior states. The most motile, most active cells in Cluster 4 are enriched for Pax7. **(D)** Representative images of Pax7/MyoG staining in cells after timelapse imaging. Panels on the far left are the final DIC image from the timelapse, registered and overlaid with fluorescent immunostains. Remaining panels are raw images prior to registration.

Clustering genes into Modules based on pseudotemporal expression patterns reveals that while many genes increase or decrease monotonically with activation as expected, other genes display non-monotonic behavior (Fig. 2F). Genes in Modules 1 and 3 display maximum expression at points in between the most quiescent and most activated states. Module 1 contains genes related to mRNA processing and splicing, as determined by gene ontology analysis (see Methods). Module 3 contains genes related to cell cycle regulation and developmental processes (Fig. S3).

These results support the notion that transcriptional programs during myogenic activation exhibit a variety of temporal behaviors, including non-monotonic and non-linear temporal regulation. Expression peaks and valleys in the nonmonotonically regulated gene modules provide evidence that there are intermediary transcriptional states of myogenic activation that are not simple interpolations of the initial and final transcriptional states.

We next identified markers of myogenic activation by differential expression between the quiescent and activated cell populations. We report differentially expressed genes that pass an effect size threshold of at least 0.15 log_2_ fold change and are expressed in at least 5% of cells in at least one side of the contrast (see Methods). Differential expression analysis revealed 3,864 genes altered by activation. Of these genes, 2,631 showed significant increases in expression while only 1,034 showed significant decreases, indicating that myogenic activation is associated with more transcriptional activation than repression. Gene ontology (GO) analysis of the differentially expressed marker genes suggests that these genes largely reflect biosynthetic and metabolic pathways, consistent with the notion that myogenic activation corresponds to a dramatic metabolic and geometric rearrangement of cellular state (Fig. S1).

This interpretation is further reinforced by weighted gene correlation network analysis (WGCNA) [33] which elucidates two gene modules during activation. The first “Quiescence Module” is upregulated in quiescent cells and contains genes related to cell stress responses, transcriptional suppression, and negative regulation of cell proliferation by GO analysis. By contrast, the “Biosynthetic Module” is upregulated during activation and contains genes related to protein biosynthesis, transcriptional upregulation, ribosome biogenesis, and RNA maturation (Fig. S4). Together, these results suggest that myogenic activation is heterogenous among individual cells at multiple time points in the process and that the activation process can be decomposed into a set of activation states reflecting cellular biosynthetic activity.

**Figure 4:**
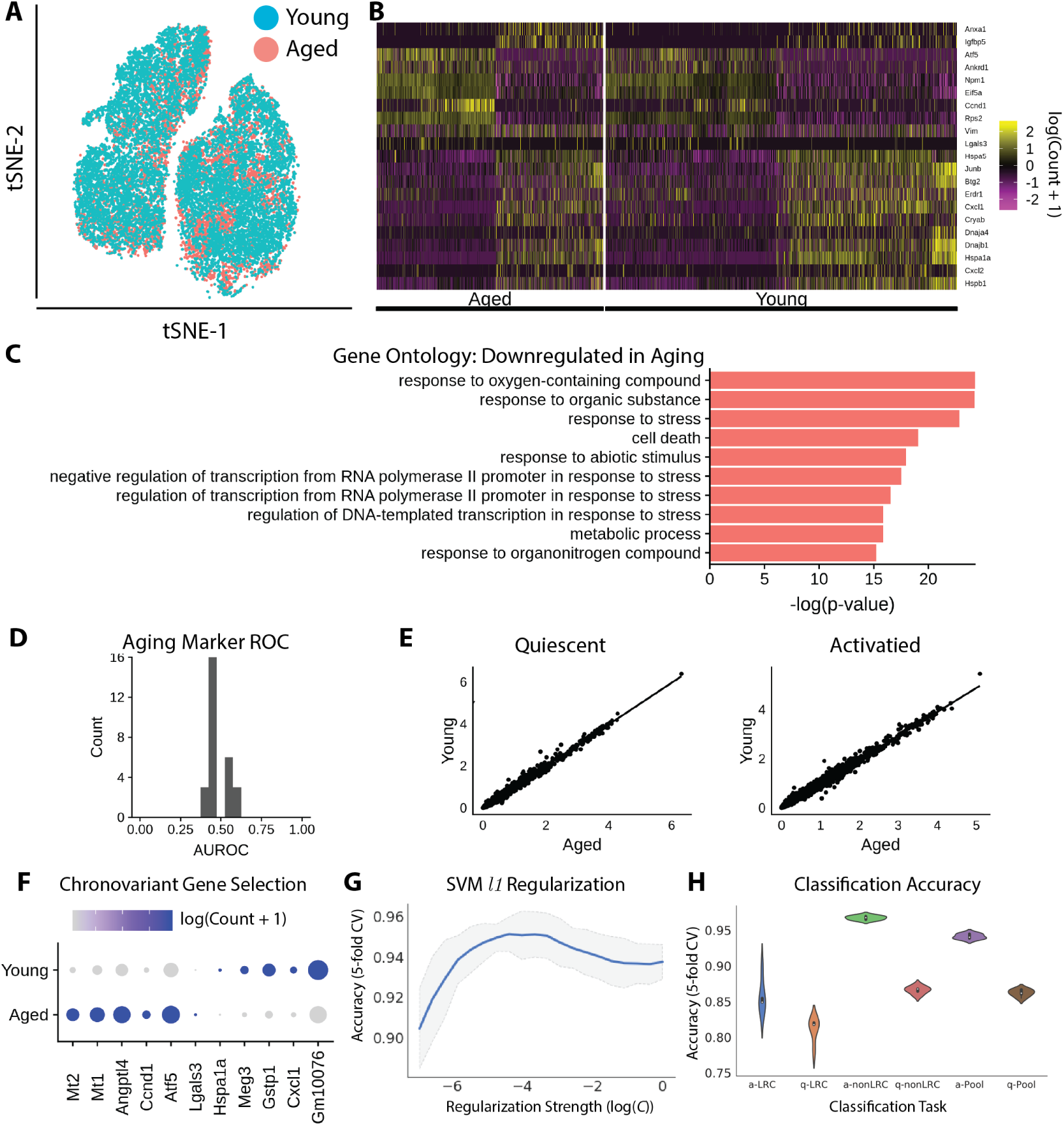
Aged MuSCs display transcriptional changes across many genes. **(A)** Aged and young cells labeled in transcriptional space (t-SNE visualization). **(B)** Heatmap of differentially expressed genes between aged and young MuSCs. **(C)** Gene ontology analysis for differentially expressed genes. **(D)** Gene-wise AUROC analysis demonstrates that single gene is predictive of MuSC age state. **(E)** Correlation of aged and young transcriptomes. **(F)** Chronovariant genes displayed on a dotplot. Darker colors indicate higher expression, larger dots indicate expression in a larger proportion of cells. **(G)** SVM classification accuracy versus regularization (*L*_1_) strength identifies a subset of genes for age discrimination. **(H)** Classification performance for aged vs. young LRCs, nonLRCs, and the total MuSC pool. All classifiers display *>* 92% accuracy.

### 2.4 Aged MuSC transcriptomes show modest differences across many transcripts

Aged MuSCs have significantly impaired regenerative capacity. We previously found that conversion of highly regenerative LRCs to less regenerative nonLRCs is a factor in this regenerative decline [11, 10]. Do aged MuSC transcriptomes reflect these functional deficits? To answer this question, we randomly sampled populations of aged and young MuSCs with physiological ratios of LRCs:nonLRCs to mimic MuSC pools *in vivo*. A young MuSC pool was sampled with a 35:65 LRC:nonLRC ratio, and an aged pool with a 15:85 ratio. These correspond to physiological ratios observed by FACS (Fig. S4).

Similar to our cell behavior analysis, we find that aged MuSCs do not segregate discretely in transcriptional space (Fig. 4A). This suggests that aged and young MuSC transcriptional states are largely overlapping. To determine whether aged cells display a preference for some states over others, we quantified occupancy of our transcriptional clusters above for both young and aged cells. There is no apparent state preference in aged cells among the transcriptional clusters (Fig. S4). This differs from the state preference of aged cells among the behavioral clusters we identify (Fig. 1D), suggesting that either the state preference arises after the 18 hour time point captured by scRNA-seq, or that the state preference is less dramatic at the transcriptional level.

Performing differential expression analysis, we identify 174 differentially expressed genes between aged and young cells when comparing both quiescent and activated cells (Fig. 4B). GO analysis of these genes indicates that they largely represent biosynthetic processes and stress responses, with protein translation processes upregulated with aging and stress responses downregulated (Fig. 4C). Among the differentially expressed genes, young cells display elevated levels of stress response heat-shock proteins *Hspb1* and *Hspa5*, suggesting that aged MuSCs are less able to mount appropriate stress responses.

In a more specific analysis, we also find differentially expressed genes with aging specifically within the quiescent and activated states. In quiescence, we find 200 differentially expressed genes between aged and young MuSCs. GO analysis suggests these genes are related to protein folding and cellular stress responses, both downregulated in aged cells (Fig. S4). During activation, we identify 275 differentially expressed genes, suggesting that activation accentuates age-related transcriptional differences. GO analysis likewise identifies that these genes are enriched for catabolic processes, downregulated in aging, and stress responses, upregulated in aging (Fig. S4). Prominently, two of the top differentially expressed genes are metallothionein proteins *Mt1* and *Mt2*, which are upregulated in aged cells. Metallothioneins have anti-oxidant and anti-apoptotic effects and are associated with increases in longevity [32, 50]. Upregulated levels in aged cells may indicate a compensatory response to age-related oxidative stress.

### 2.5 Gene expression variance is altered by aging

In addition to information about mean gene expression levels, single cell RNA-seq provides information about the variation in expression within cell populations. Recent work has suggested that aging may increase gene expression variance in immune cells [35]. Does a similar increase in gene expression variance occur in aged MuSCs? To answer this question, we employ the difference from the median (DM) method to quantify the amount of “overdispersion,” or variance above expectation, for each gene [29]. This “overdispersion,” metric is necessary due to the confounding relationship between mean expression levels and measured cell-to-cell variation.

Measuring overdispersion, we find that aged cells have higher dispersion in the quiescent state (*p <* 0.05, Wilcoxon Rank Sums). Surprisingly, this difference in variability reverses in activated cells, such that activated young cells have higher overdispersion than activated aged cells. (*p <* 0.001). Additionally, young cells show increased gene expression variance as a result of activation, while aged cells show decreased gene expression variance upon activation (*p <* 0.001, both instances)(Fig. S5).

**Figure 5:**
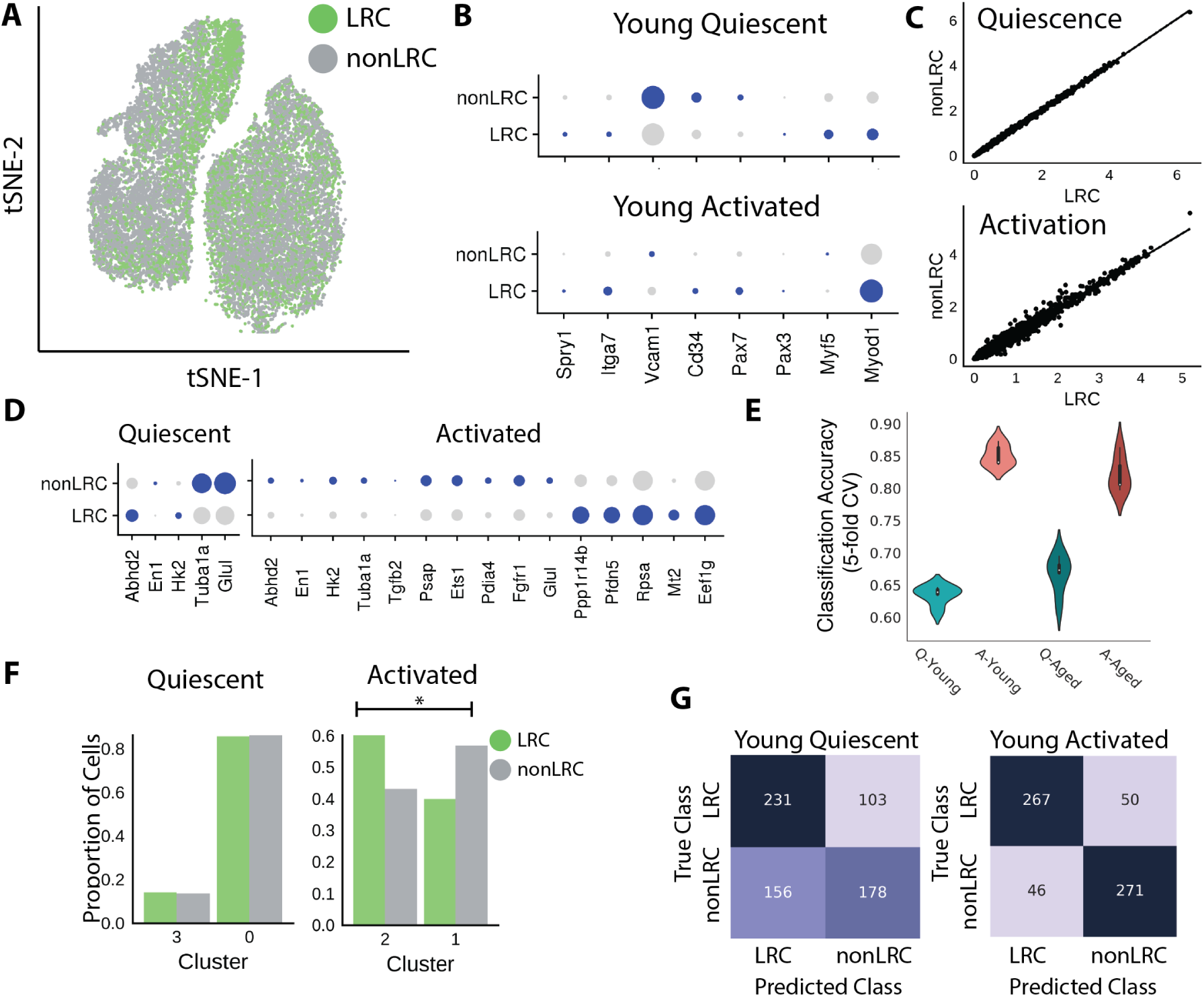
Activation induces differential responses in LRCs and nonLRCs. **(A)** LRCs and nonLRCs labeled in transcriptional space (t-SNE visualization). **(B)** Expression of known myogenic regulatory genes in quiescent and activated LRCs and nonLRCs. Larger dots indicate a greater proportion of expressing cells, darker colors indicate higher expression. **(C)** Correlation between LRC and nonLRC mean gene expression values in quiescence and activation. **(D)** Differentially expressed genes between LRCs and nonLRCs in quiescent and activated conditions. Few transcriptional differences are present in quiescence, but activation manifests differential expression across many genes. **(E)** LRC:nonLRC classification performance across activation states and ages. Classifiers more readily discriminate LRCs from nonLRCs in activation. **(F)** Distribution of quiescent and activated LRCs/nonLRCs in transcriptional clusters indicates a heterogeneous activation response. **(G)** Confusion matrices for LRC:nonLRC classifiers. Confusion is higher in the quiescent condition.

This result suggests that the levels of gene expression variance are altered by aging in a context-dependent manner. Importantly, it is possible that the decreased gene expression variance we find in activated, aged MuSCs is due to selective pressure during the *in vitro* activation process. If activation selected for a more stringent subset of cells in aged MuSCs than young counterparts, we may observe less variance in the aged cells as a result of that selection.

### 2.6 Discrimination of aged and young MuSCs by machine learning

The field of aging biology seeks biomarkers of aging that can be used as an assay of the aged phenotype. To determine if any genes would serve as effective biomarkers at the transcriptional level, we performed receiver operator characteristic (ROC) analysis for each differentially expressed gene between young and aged cells. No gene provides an area under the ROC (AUROC) greater than roughly 0.6, suggesting that no single gene acts as an effective biomarker for aging (Fig. 4D).

Can we identify a predictive relationship between multiple genes and MuSC age? Predicting a categorical response such as age from a set of many continuous descriptors is a classical machine learning classification problem. We developed a support vector machine (SVM) classifier approach to classify aged and young MuSCs by incorporating information from multiple differentially expressed genes. Briefly, we identified a set of candidate “chrono-variant” genes that change with age, focusing on the case of activated MuSCs where differences are more pronounced (Fig. 4F).

We trained linear SVM models with *L*_1_ regularization to enforce sparsity within the provided gene set, identifying a smaller subset of genes from which to predict MuSC age. The regularization strength *C* for our models is optimized using 5-fold cross-validation on a training data set (Fig. 4G). For model validation, we utilize a “hold-out” set containing a random sample of 20% of the total data set. This data set is placed behind a “firewall” and is not used in model construction or optimization in any form.

We first applied this machine learning approach to predict age within LRC and nonLRC subsets. We also separately classified quiescent and activated time points. This analysis controls for the difference in LRC:nonLRC ratio between aged and young MuSC pools. Classifying activated LRC age, we identify a set of 54 genes that yield a prediction accuracy of roughly 85% (hold-out validation) (Fig. 4H). For nonLRC classification, we identify 104 genes that yield a predictive accuracy of roughly 96% (hold-out validation) (Fig. 4H). Of these genes, only 40 are common to both LRCs and nonLRCs, suggesting that transcriptional aging manifests differently in LRC and nonLRC populations. Classifying the activated young and aged MuSC pools (each sampled to model physiological LRC:nonLRC ratios), we identify a set 99 genes that provide roughly 95% classification accuracy (Fig. 4H)(Fig. S5). In each case (LRC, nonLRC, pooled), we find that classification of activated cells is more effective than classification of quiescent cells, further suggesting that activation reveals transcriptional aging phenotypes. This classification result represents the first effective assay to discriminate the age of individual muscle stem cells.

### 2.7 Estimating the contribution of LRC to nonLRC conversion to transcriptional aging

The proportion of LRCs in the MuSC pool is roughly 35% in young animals and decreases to roughly 15% with age. How much does this conversion of LRCs to nonLRCs with age contribute to the overall transcriptional changes we see with aging? By fitting probabilistic classification models, we can use the ability of a classifier to discriminate between young and aged MuSC populations as a metric of the “magnitude” of transcriptional change (see Methods). This technique, known as the density ratio trick, is often used in machine learning to estimate the difference between two distributions [49].

Populations of aged and young cells were simulated by random sampling with either physiologically observed LRC:nonLRC ratios or equal LRC:nonLRC ratios (both aged and young sampled with 35:65 LRC:nonLRC ratios). Fully-connected neural network models were trained to discriminate aged and young cells using a softmax layer for probabilistic output (see Methods). Classifier accuracies are not notably changed when the LRC:nonLRC ratio is changed from equal to physiologically observed ratios. Likewise, the divergence metric we compute from the probabilities output by the classifiers is comparable in both conditions. The similarity in classification accuracy and divergence magnitude in the face of changes to the LRC:nonLRC ratio suggests that LRC to nonLRC conversion does not dramatically alter the “magnitude” of age-related transcriptional change (Fig. S6).

**Figure 6:**
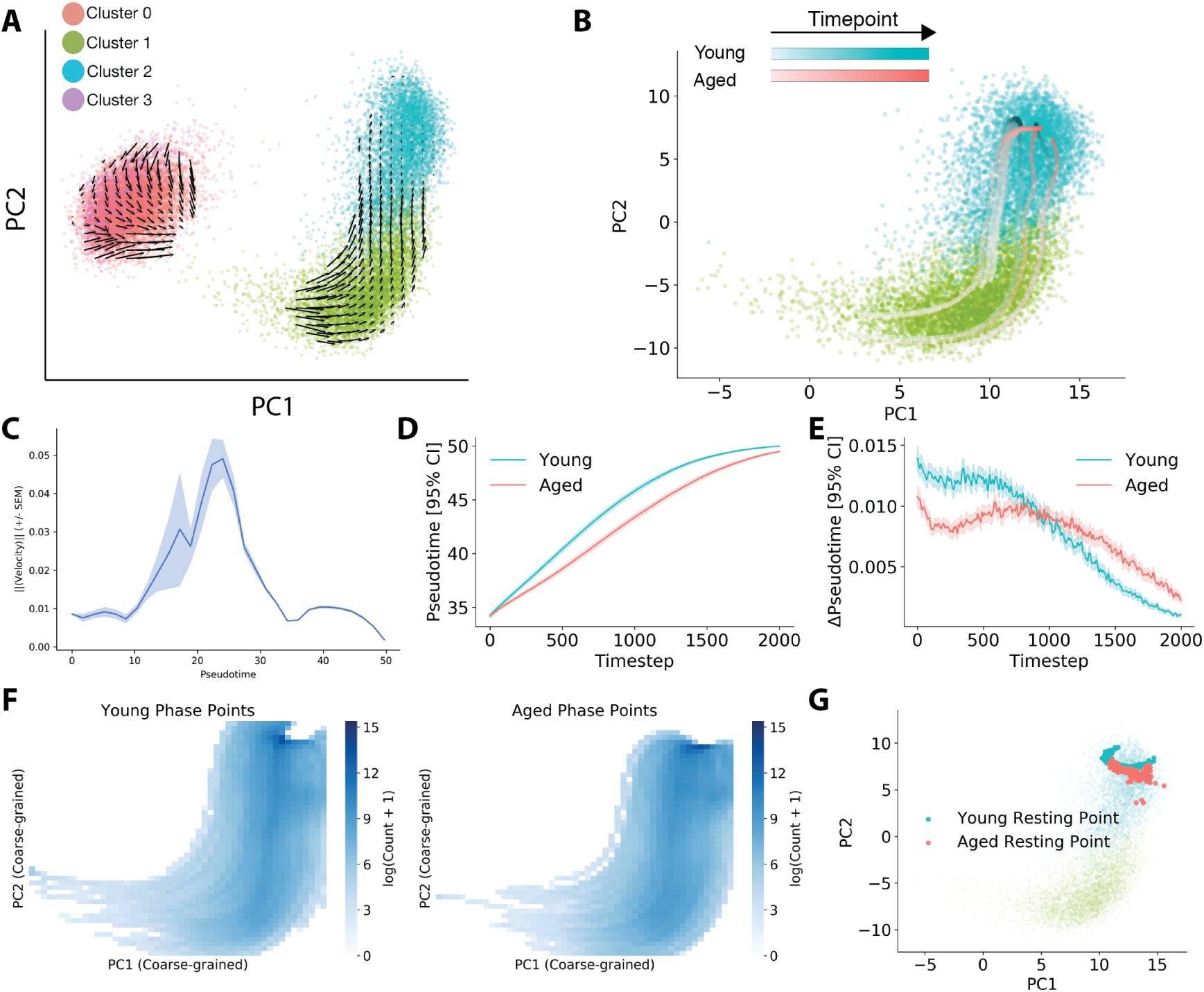
Aged MuSCs transition aberrantly through transcriptional space. **(A)** MuSC transcriptional space overlaid with arrows representing the direction and magnitude of RNA velocity at each state location. Colors indicate transcriptional state identity. **(B)** Representative phase point simulations in aged and young RNA velocity fields, overlaid on the activated MuSC cells in a PCA embedding. **(C)** State transition rates as measured by RNA velocity magnitude across pseudotime using a rolling mean. **(D)** Predicted pseudotime progression for phase point simulations in either aged (red) or young (blue) velocity fields. **(E)** Change in pseudotime for phase point simulations at each timestep. **(F)** Heatmap representing the mean density of phase points at each point in PC space across the entire simulation. Young and aged phase simulations show qualitatively similar trajectories through state space. **(G)** Terminal locations of young and aged phase point simulations in PC space, overlaid on cell locations. Both young and aged simulations show similar final resting positions.

### 2.8 Activation manifests transcriptional differences due to proliferative history

LRCs are functionally distinct from nonLRCs, and this functional difference persists throughout life [10, 11]. While we find that the majority of transcriptional differences between aged and young cells are independent of proliferative history, we also investigated whether LRCs and nonLRCs were transcriptionally distinct from one another. Similar to aged and young cells, LRCs and nonLRCs appear to share a transcriptional state space, and do not readily segregate along an axis in reduced dimensional space (Fig. 5A). Likewise, the mean gene expression levels between the two states have a near perfect correlation in quiescent cells (*r* = 0.99, Fig. 5C).

To determine what transcriptional differences underlie the LRC/nonLRC functional differences in an unbiased manner, we examined differentially expressed genes between the two pools in young MuSCs, so as to avoid confounding effects of aging. Differential expression analysis reveals 97 differential genes between the LRC and nonLRC pools.

Considering quiescent and activated cells separately, there are only 14 differentially expressed genes in quiescent cells, but 195 differentially expressed genes in activated cells (Fig. 5D). Differentially expressed genes include known stress response genes such as heat shock proteins and inflammation signatures, largely enriched within the nonLRC population. Resiliency and biosynthetic genes are also differentially expressed, with known longevity factor [50] *Mt2* and protein translation components *Pfdn5* and *Eef1g* enriched in LRCs. GO terms for nucleoside triphosphate metabolic processes are enriched in genes upregulated in young LRCs, while terms for cell death and apoptotic processes are enriched in young nonLRC upregulated genes. These expression differences suggest that LRCs may possess a biosynthetic advantage over nonLRCs, while nonLRCs exhibit a pronounced stress response. This is consistent with the recent identification of a stress-tolerant subset of adult LRCs [45].

Using biological priors to guide analysis, we investigated expression of known regulators and markers of myogenic state in LRCs and nonLRCs. The data reveal an activation dependence on the relative enrichment of multiple myogenic markers. *Pax7* and *Cd34* are slightly enriched in nonLRCs in quiescence, but enriched in LRCs in activation. Similarly, LRCs are shown to be enriched in *Vcam1* in quiescence, but express less of this marker in activation (Fig. 5B). This finding is again consistent with a differential response to activation in LRC and nonLRC states, as suggested by mean expression value correlations and unbiased differential expression analysis. Considering the distribution of LRCs and nonLRCs among clusters of activation further confirms a heterogeneous activation response. In the quiescent state, LRCs and nonLRCs do not display significant differences in distribution between the two quiescent transcriptional clusters. However, after activation, LRCs are significantly enriched in the most activated transcriptional Cluster 1. Roughly 60% of activated LRCs are in the most activated cluster, compared to only 40% of the activated nonLRCs (Fig. 5F). The higher proportion of LRCs in the most activated region of transcriptional space suggests that LRCs may activate more quickly than nonLRCs.

Differential expression analysis also suggests a heterogeneous activation response. Regression analysis of mean gene expression levels between the LRC and nonLRC states corroborates this finding. The two states have a near perfect correlation in quiescent cells. In activation, the mean gene expression values are less correlated (*p <* 0.001, Fisher’s *r* to *z* transformation, Fig. 5E), indicating that activation induces different transcriptional responses in LRCs and nonLRCs.

This result is surprising, as LRCs are believed to be “reserve” stem cells, while nonLRCs are presumed to be precocious in their response based on differentiation assays [10, 11]. To confirm this observation with an orthogonal assay, we measured EdU incorporation in young LRCs and nonLRCs during activation in culture. MuSCs were cultured for 50 hours, and pulsed with EdU 12 hours and 2 hours prior to fixation. LRCs incorporated EdU at a higher rate (21%) than nonLRCs (17%) across three animals and *n* ≈ 20, 000 cells (*p <* 0.001*, χ*^2^*test*)(Fig. S6). Collectively, these results suggest that LRCs activate more rapidly than nonLRCs, based both on progression through transcriptional space and the timing of cell cycle entry.

Similar to our classification of MuSCs from different ages above, we trained similar classifiers to discriminate LRCs and nonLRCs. We trained a separate model to classify LRC/nonLRCs at each timepoint and each age. Classification of quiescent cells performs poorly at both ages (roughly 65% accuracy), while classification of activated cells is more effective – roughly 85% accuracy for young cells and 80% for aged cells (Fig. 5E, G). As in our age classification experiments, this result suggests that activation reveals differences between cell populations that are masked in quiescence. Regularization with an *l*_1_ penalty identifies 102 genes that optimize LRC/nonLRC classification of young activated cells, and 72 genes that optimize classification of aged activated cells. 35 genes are shared between these sets, enriched for apoptotic and oxidative stress response gene sets, suggesting that differences in the stress response of LRCs and nonLRCs may be some of the clearest distinguishing features across ages.

### 2.9 Transcriptional kinetics are aberrant in aged MuSCs

The lack of unique aged transcriptional states and modest differential expression results between aged and young cells are surprising in light of the dramatic differences in functional potential between aged and young cell populations [11, 15]. These results suggested to us that the rate at which aged and young cells activate may be an additional source of variation that contributes to their functional differences. To quantify rates of phenotypic change between the MuSC transcriptional states during activation, we utilized the recently developed RNA velocity method [31]. This method estimates a “velocity” of transcription, or rate of change in the transcript level, by estimating the decay rates of measured, fully spliced mRNAs, and estimates the rate of mRNA transcription using ratios of spliced to unspliced reads.

Performing RNA velocity estimation on all 20, 000+ single MuSC transcriptomes shows that each state of MuSC transcription gives rise to a neighboring state in the sequence inferred by pseudotiming (Fig. 6A). As an internal validation check, we find that RNA velocity indeed indicates that quiescent cells (sequenced immediately after isolation) are moving toward activated cells (sequenced after 18 hours in culture) in transcriptional space. This result provides further confirmation that the ordering of transcriptional clusters we infer by pseudotiming is correct.

The magnitude of mean RNA velocity represents the rate of collective phenotypic change at the transcriptional level for a given group of cells. This approach provides an inferred measurement of state transition rates in transcriptional space, similar to the measurement of state transition rates we make by direct observation in cell behavior space. Quantifying the magnitude of RNA velocity across pseudotime in MuSCs reveals that RNA velocity follows a concave curve (Fig. 6C). Concave transition rates across pseudotime suggest that myogenic activation is a switch-like process, corroborating our earlier observations made by cell behavior analysis [28]. Consistency in state transition measurements between RNA velocity and cell behavior phenotyping suggests that cell behavior state transitions reflect the underlying transcription state kinetics.

Do aged and young MuSCs move through transcriptional state space differently? To answer this question, we developed a method to model cellular progression through transcriptional space using phase point simulations. RNA velocity generates a vector field in transcriptional state space. Phase point analysis is classic dynamical systems method to investigate the properties of a vector field where a simulation is performed to determine how a particle might flow along a vector field, as if it were a floating leaf carried by currents in a river [48]. Here, we simulate a set of phase points that begin in the more primitive regions of transcriptional space occupied by cells from our “activated” 18 hour timepoint and evolve them over time using velocities inferred from either young or aged cells nearby in transcriptional space (see Methods). Given these simulated trajectories through transcriptional space, we ask whether notable differences are present in phase points simulated using young velocities (“young phase points”) relative to those simulated using aged velocities (“aged phase points”).

One question is whether aged and young phase points progress through the process of activation at different rates. To assess progress through cell activation, we trained a *k*-nearest neighbors regression model (kNN-R) to map transcriptome PCA embeddings to pseudotime coordinates as determined using Monocle 2 (see Methods, Fig. S5). Scanning a range of parameters, we find a regressor using *k* = 30 nearest neighbors is sufficient to achieve *r*^2^0.96 predicting pseudotime values from the first two principal components (Fig. S6, 5-fold cross-validated).

For each timestep in a phase point simulation, we predict the pseudotime coordinate of the point using this model. Comparing inferred pseudotime coordinates for young phase point simulations and aged phase point simulations, we find that young phase points progress more rapidly through the activation process than aged phase points (Fig. 6D). Computing numerical derivatives for pseudotime coordinates ΔPseudotime, young phase points appear to progress more rapidly from the earliest time steps (Fig. 6E). This result suggests that aged cells may progress more slowly than young cells through the activation process in a similar manner to the phase point simulations. Repeating these analysis using age-dependent initialization points and/or using noiseless trajectory simulations yields similar results (Fig. S7).

**Figure 7:**
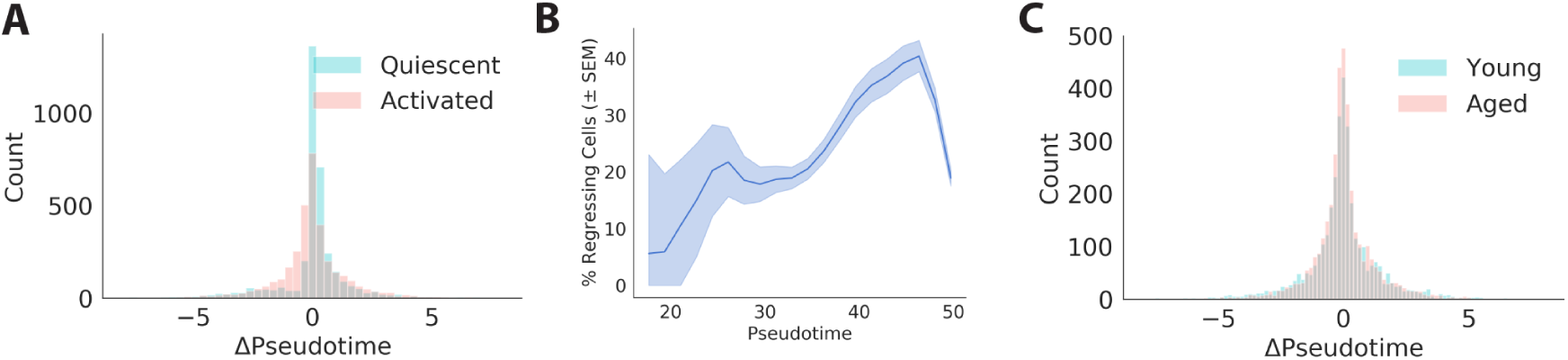
MuSCs exhibit lineage regression during activation. **(A)** Distribution of changes in pseudotime (ΔPseudotime) inferred from RNA velocity estimates in young quiescent and activated cells. Activated cells regress more frequently. **(B)** Proportion of cells moving backward in pseudotime as a function of position along the pseudotime curve for cells in the later experimental timepoint (18h *in vitro*). **(C)** Distribution of changes in pseudotime for young and aged MuSCs across conditions. Aging does not appear to dramatically influence lineage regression frequency.

Phase point simulations additionally provide information about the location of “attractors” in transcriptional space. Attractors are locations in a state space where phase points tend to converge and come to rest. By measuring the density of phase points in transcriptional space and examining the positions where they come to rest, we can identify attractors and determine if young and aged phase point simulations share attractors, or have unique attractors.

Visualizing the density of phase points in transcriptional space as the total number of phase points to pass through a region, there are qualitatively few differences in the shape of trajectories between young and aged cells (Fig. 6F). Focusing on the locations where phase points come to rest, there are likewise modest differences in the specific shapes of attractor states, but overall similar attractor positions between young and aged phase point simulations (Fig. 6G). Collectively, results of these phase point simulations suggest that the set of intermediate transcriptional states a MuSC visits in the course of activation is largely similar between young and aged cells. However, aged cells appear to activate at a slower rate than young counterparts. Each of these points supports the Different Rates model of aging pathology outlined above.

### 2.10 Lineage regression occurs during myogenic activation in a subset of MuSCs

The discovery of reserve cells generated during myogenic commitment more than 20 years ago first presented the idea that MuSCs may revert to earlier stages in the lineage progression under some conditions [57]. It is currently unclear how frequently MuSCs transition “backwards,” in the myogenic activation program. We assessed the frequency of MuSCs transitioning backward in the lineage progression by quantifying a “change in pseudotime” (ΔPseudotime) for each cell in our young MuSC single cell RNA-seq data set. ΔPseudotime was estimated using the *k*-nearest neighbors regression (kNN-R) model referenced above. Pseudotime values were predicted for the “future” transcriptomes inferred by RNA velocity, and the difference between the predicted pseuodotime and observed pseudotime values was taken as the ΔPseudotime (see Methods for details). We defined a cell as “regressing” in pseudotime if ΔPseudotime was more than 1/2 standard deviation below 0.

This analysis reveals that roughly 16% of young MuSCs are regressing in pseudotime during the period of myogenic activation we measure (Fig. 7A). Regression is more frequent in activated (≈ 20%) than quiescent MuSCs (≈ 15%). Intriguingly, this fraction of cells is similar to the fraction of cells which regress into the “reserve cell” state. Quantifying the frequency of “lineage regression” across pseudotime for cells from our later timepoint (18 hours *in vitro*) reveals that cells regress more frequently in the later stages of activation we observe (Fig. 7B). This regression behavior appears robust to age-related changes (Fig. S6). These results suggest myogenic activation is a two-way process even under growth-promoting conditions, perhaps resembling a biased random walk in transcriptional space.

## 3 Discussion

Dynamic changes in stem cell phenotypes are essential for both development and regeneration. However, due to the difficulty of measuring single cells over time, quantitative understanding of these processes remains elusive [1]. Changes in these cell state transition rates may explain some portion of the decreased regenerative potential observed in aged stem cells, as in the muscle stem cell system.

Some dynamic information can be inferred from static ensemble measurements, a class that includes destructive molecular assays with single cell resolution such as immunostaining or single cell transcriptomics. This type of inference is unable to assess if the progression of cells through a set of states is uniform or heterogeneous, if intermediary states are bistable or transient, or to determine if a given cellular feature influences velocity [55]. Answering either of these questions requires some additional dynamical information – essentially, individual cells must be measured at multiple time points.

Here, we use timelapse imaging and single cell RNA-sequencing to provide paired measurements of this nature to investigate muscle stem cell activation. Previous work has identified “primed” states of activation, but it is unknown whether these states are bistable or transient [40, 41]. If a primed state was bistable, we would expect to observe cells entering it from both the “less activated” end moving forwards, and also the “more activated” end moving backwards. Using paired measurements, we do not find evidence of bistable states within a continuous activation process on the timescales we observe. Our RNA velocity analysis does not identify any intermediate basins of attraction as indicated by the smooth sigmoidal ΔPseudotime curves (Fig. 6). In this analysis, an additional attractor state might appear as plateau on this activation curve. This suggests that primed cell states are transient – akin to an “out-and-back” journey down the path of activation.

Static ensemble read-outs during MuSC activation have long demonstrated that MuSCs occupy different transcription factor states, even at a single time point [14]. However, these measurements could not explain where in the activation process heterogeneity arose. Here, we find that MuSCs progress through the activation process stochastically, with a non-trivial proportion of the population moving “backwards” through the activation process. This suggests that the heterogeneity of MuSC positions along the activation trajectory arises as an accumulation of differences in the rate of cell state transitions. These differences appear to be both stochastic and associated with distinctive features between MuSC subpopulations. Although the macroscopic processes of muscle development and regeneration proceed without these apparent reversals, these observations indicate that phenotypic change at the cellular level may involve considerably more noise. This is reminiscent of the qualitative differences between the physical motion of macroscopic objects, like a ball rolling down a hill, and microscopic motion, where noise can dominate the movement of small molecules which often reverse direction completely.

Aging leads to dramatic declines in the regenerative capacity of MuSCs. By single cell RNA-sequencing, we surprisingly find minor transcriptional differences between aged and young cells. Notably, these differences are more pronounced in activated cells than quiescent cells, suggesting that the functional challenge of an activation stimulus (in this case, cell culture) manifests age-related changes at the transcriptional level that are latent in quiescence. Agerelated changes may be similarly hidden from common transcriptional assays in the absence of functional challenge, as observed by others [27, 35, 15, 54]. The large degree of similarity between transcriptional states of aged and young cells over the course of activation suggests that the sequence of cellular states, or trajectory of activation, is preserved with aging.

Measuring state transition rates during activation reveals that aged MuSC have dampened state transition rates. By behavioral analysis, aged MuSCs display a preference for less motile, less activated states and decreased rates of transition into more active states. Similarly, phase point analysis of RNA velocity vectors suggests that aged cells transition more slowly through transcriptional states during the earliest phases of activation than young counterparts. These data support a conceptual model in which aging MuSCs exhibit “Different Rates” of activation, even if they follow the same trajectory. Measurement of cell state dynamics in other stem cell pools may reveal if dampened cell state transition rates are a common feature of stem cell aging.

## 4 Methods and Materials

### 4.1 Animals

Animals were handled according to UCSF Institutional Use and Care of Animals Committee (IUCAC) guidelines. All experimental mice were male *Mus musculus* of the C57Bl/6 background. Aged mice for cell behavior experiments were 20 months of age. Young mice for cell behavior experiments were 3-5 months of age. All mice for single cell sequencing experiments harbored *H2B-GFP +/-; rtTA +/-* allelles and were developmentally labeled for proliferative history by administration of doxycycline E10.5-E16.5. Aged mice for RNA-seq sequencing experiments were 20 months old and young mice were 3 months old. All mice were born at UCSF and aged in-house.

### 4.2 Cell Isolation and Culture

Muscle stem cells were isolated by FACS using a triple negative CD31^-^/CD45^-^/Sca1^-^ and double positive VCAM^+^/*α*7-integrin^+^ strategy as described [11]. Antibodies are from the following suppliers: PE-Cy7 Rat anti mouse CD31 Clone 390, PE-Cy7 Rat anti mouse CD45 Clone 30-F11, and APC-Cy7 Rat anti mouse Ly-6A/E Clone D7 (all BD Pharmigen); Mouse CD106/VCAM1 PE (Invitrogen).

Cells for behavior analysis were seeded at 850 cells/well on sarcoma-derived ECM (Sigma, St. Louis, MO) in 96-well plates. Cells were maintained in rich growth media (F10 (Gibco), 20% FBS (Gibco), [5 ng/mL] FGF2 (R & D)). For single cell sequencing experiments where cells were activated, cells were seeded in sarcoma-derived ECM coated 6-well plates and allowed to activate in plating media (DMEM, 10% horse serum) for 18 hours prior to library preparation. For each behavior experiment, 1 young (3 months old) and 1 aged mouse (20 months old) were used as sources of young and aged MuSCs, respectively.

### 4.3 Timelapse Imaging and Cell Behavior Analysis

MuSCs were imaged in 96-well plates on an incubated microscopy platform (Oko Lab) for 48 hours. Images were collected with DIC contrast every 6.5 minutes to track cell movement. Thirty rasterized fields-of-view at 20X magnification were collected from each well using an Andor Zyla 4.2 camera with pixel size of 6.5 *µ*m. Images were segmented using a fully-convolutional DenseNet-103 neural network model, following the architecture of [25]. The model was implemented in PyTorch and trained on manually segmented images from each experiment. Code for our implementation is available at https://github.com/jacobkimmel/fcdensenet pytorch. Cell tracking was performed using a custom bipartite tracking implementation that utilizes a Kalman filter motion model. Python code for our tracking implementation is available at https://github.com/jacobkimmel/musc tracker. The first 15 hours of each movie were not analyzed due to mechanical jitter present while culture plates settle into position in the microscopy rig. Cell behavior was analyzed using *Heteromotility*, as previously described [28] and available at https://github.com/cellgeometry/heteromotility. GNU parallel was used to parallelize multiple portions of the analysis [51].

For paired behavior-immunocytochemistry experiments, cells were fixed in 4% paraformaldehyde for 10 minutes immediately following the imaging timecourse. All steps were carried out at room temperature, unless otherwise noted. Cells were washed in PBS 3X, using gentle pipette aspiration (without vacuum) to remove buffer. We found that vacuum aspiration tends to dislodge a large number of cells. Cells were subsequently permeabilized with 0.2% PBSX (PBS + Triton X-100) in two 5 minute washes. After permeabilization, cells were blocked in 10% goat serum in PBSX for 60 minutes. We added primary antibodies for Pax7 (Mouse, Developmental Studies Hybridoma Bank) and MyoG (Rabbit, Santa Cruz Biotechnology Cat sc-576 AB 2148908) at 1:100 concentrations in 10% goat serum/PBSX overnight at 4*^o^*C. Cells were washed 4X in PBSX, then blocked a second time by incubation in 10% goat serum/PBSX for 60 minutes. Cells were incubated with secondary antibodies anti-mouse Alexa 488 (Thermo Fischer) and anti-rabbit Alexa 647 (Thermo Fischer) for one hour. Cells were finally washed 3X in PBSX, 5 min each, then 3X in PBS, 5 min each, incubated with Hoescht 33342 (5 *µ*g/mL in PBS) for 10 minutes, and washed with PBS again.

After staining, cells were returned to the same timelapse microscopy system used for behavioral imaging, and fluorescent images were captured. We performed segmentation of the final brightfield image from the behavioral timecourse, and the nuclear channel of the fluorescent stained image, then performed image registration using nearest neighbors to match immunofluorescence signals to cell behavior tracks.

### 4.4 EdU Staining

Muscle stem cells were isolated as described above from *n* = 3 *H2B-GFP +/-; rtTA +/-* mice (4 months old, labeled developmentally as above) and cultured in plating media (DMEM, 10% horse serum) for 50 hours. EdU was pulsed into the media at 12 hours and 2 hours (10 *µ*M final concentration) prior to fixation with 4% PFA for 15 minutes. Staining followed the Click-iT EdU Alexa Fluor 647 kit protocol (Thermo-Fisher).

### 4.5 Single cell RNA-sequencing

MuSCs were isolated from 2 young (3 m.o.) and 2 aged (18 m.o.) male *H2B-GFP; rtTA* heterozygous mice. Mice were all labeled to capture proliferative history during development by doxycycline induction of *H2B-GFP*. This allowed for isolation of label retaining MuSCs (LRCs) and and non-label retaining MuSCs (nonLRCs) based on GFP intensity, as described [10]. Half of collected cells were immediately transferred to a 10X Genomics Chromium system for library preparation using the 10X 3’ Single Cell v2 chemistry. The remaining cells were activated by culture on ECM-coated cell culture dishes for 18 hours in plating media (10% horse serum, DMEM), then dissociated using Cell Dissociation Buffer, stained with PI, and sorted by FACS to remove dead cells (PI negative). Live, activated cells were transferred to the 10X Chromium system for identical library preparation. Libraries were pooled and sequenced using an Illumina NovaSeq platform.

### 4.6 Single Cell Transcriptome Analysis

Raw sequencing data was demultiplexed using Illumina bcl2fastq. Demultiplexed sequencing reads were aligned to the mouse transcriptome using the STAR aligner [17]. Individual UMIs were detected and assigned to corresponding cell barcodes using 10X Genomics *cellranger*, samplewise. Droplets containing cells were identified using a heuristic implemented in *cellranger* to call a threshold on a plot of Barcodes vs. Number of Assigned UMIs. Individual libraries were aggregated using *cellranger* and normalized to the same sequencing depth by random sampling of reads. Libraries were quality controlled by examining Sequencing Depth vs. Unique UMI plots.

A Genes × Cells count matrix was generated from the aggregated libraries. Suspected dead cells were removed if a high proportion of total UMIs in the cell mapped to mitochondrial genes [24]. Putative doublets were removed as outliers on a histogram of UMIs/Cell and Genes/Cell [9]. Prior to normalization, the annotated transcripts *Gm42418* and *AY036118* were removed from the count matrix. These transcripts overlap an unannotated *Rn45s* rRNA locus, and may include counts from rRNA molecules that were amplified during library preparation despite polyA-selection.

Raw counts were log normalized using the Seurat [44] “NormalizeData” function, which performs

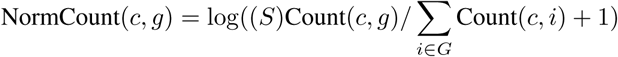

where NormCount(*c, g*) is the normalized count output for gene *g* in cell *c*, Count is the matrix of raw UMI counts, *G* is the set of all genes detected in the experiment, and *S* is a scaling factor set to 10, 000. A scaled set of counts was generated using the “ScaleData” function in Seurat which centers each normalized gene to a mean expression of *µ* = 0 and scaled the standard deviation to *σ* = 1. Variable genes were identify using “FindVariableGenes” in Seurat, and principal component analysis (PCA) was performed on the variable gene set. *t*-SNE was performed on the principal components with perplexity *p* = 30.

### 4.7 Contribution of factors to transcriptional variation

The proportion of variation explained by each experimental factor in our multi-factor experiment was estimated following the approach of Robinson et. al. [39]. Linear models were fit for each gene in the form:

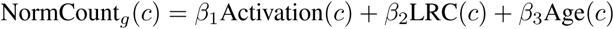

where Activation, LRC, and Age are binary vectors indicating the activation status, LRC status, and age of each cell, *c* is a cellular index, and *g* is a gene index. The proportion of variance attributable to each of these factors was calculated using an analysis of variance (ANOVA).

### 4.8 Overdispersion Analysis

Overdispersion scores were computed using the difference from the median (DM) method [29]. We eliminate all genes with a mean expression lower than *µ* = 0.1, as technical noise for genes with very low mean expression is known to be high [29].

We define an overdispersion score DM as:

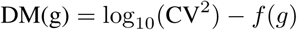

where *g* is a gene in the set of analyzed genes *G* and *f* (*·*) is a rolling median. We compute the rolling median on the mean expression (log_10_(*µ*)) vs. coefficient of variation squared (log_10_ CV^2^) plot using a bin size of *n* = 50 genes and a stride of *s* = 1 along the mean expression axis and calculating the median log_10_ CV^2^ of each bin. An additional parameter *α* was computed as the proportion of cells expressing a given gene (elsewhere referred to as the “proportion of non-zero cells”).

### 4.9 Estimation of LRC to nonLRC contribution to transcriptional change

We estimate the “magnitude” of transcriptional change with aging between a set of young and aged transcriptomes by training probabalistic classifiers to estimate the density ratio between distributions of young and aged transcriptomes. This method is commonly employed in machine learning and is known as the “Density Ratio Trick” [49, 42].

We generate populations of aged and young MuSCs by random sampling with *n* = 1000 cells per age. We sample populations with either physiologically observed LRC:nonLRC ratios (35:65 young, 15:85 aged) or equal ratios for both ages (35:65 young, 35:65 aged). The latter sampling scheme simulates a condition where LRC proportions do not change with age. Fully-connected neural networks with 3 hidden layers, each containing 100 hidden units are trained to output probabilities that a given transcriptome is either young or aged using a softmax activation. Networks are trained using a crossentropy objective and the Adam optimizer with the *scikit-learn* implementations. Networks were trained for a maximum of 1000 epochs using early stopping with a patience of 50 epochs using 10% of training data as test data for model selection. Minibatch sizes of 128 transcriptomes were used. Training was performed using 5-fold cross-validation, such that each predicted probability for a given cell was produced using a classifier that did not see that cell during training. All training parameters were chosen empirically without hyperparameter optimization.

Once trained, these probabilistic classifiers output a probability *p*(*x*) that a given cell *x* comes from the distribution of young transcriptomes, as well as a probability *q*(*x*) = 1 – *p*(*x*) that the cell comes from the distribution of young transcriptomes. The “density ratio” for each cell is simply the ratio of these two probability distributions.

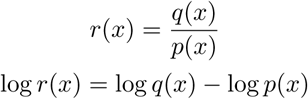

From this ratio, the Kullback-Leibler (KL) divergence can be estimated:

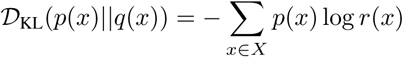

We use this estimate of the KL divergence as a measure of the magnitude of difference between young and aged transcriptomes. Because the KL divergence is asymmetric, we present the divergence measures for both directions 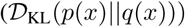 and 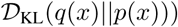. An estimate of the divergence obtained simply due to cell-cell variation within each age is computed by training classifiers on two random samples of young cells or two random samples of aged cells, where classification should perform poorly as both samples are drawn from the same distribution. As expected, our estimate of the KL divergence from these null classifiers is significantly lower than the estimates we find in classifiers trained to discriminate young from aged random samples with either physiological or equal LRC:nonLRC ratios.

### 4.10 Pseudotiming

Pseudotime analysis was performed using the Monocle2 package [38]. Genes for pseudotemporal ordering were determined by differential expression analysis between the transcriptional clusters, as described in the following section. Pseudotiming was performed on all 20,000+ cells that passed QC simultaneously in the same transcriptional state space utilizing the DDRTree method with 2 components.

### 4.11 Differential Expression Analysis

Differentially expressed genes between two populations *A* and *B* were determined based on a significant difference detected by the Wilcox Rank Sum test. Genes with a fold-change less than 0.25, or genes expressed in less than 10% of cells were filtered out from the analysis to refine the gene set. Receiver operating characteristic (ROC) scores were generated for each gene *g* based on a simple binary threshold classifier trained on only the expression of gene *g*.

### 4.12 Gene Ontology and Pathway Analysis

Gene ontology enrichment analysis was performed using g:Profiler and the *gProfileR* package, considering enriched GO terms for biological processes and KEGG pathways.

### 4.13 Support Vector Machine Classification

Support vector machine (SVM) classification models to discriminate cell age and LRC status were trained using **scikitlearn** implementations [37]. For age classification, activated MuSCs were subsampled to match the physiologically observed LRC:nonLRC ratio across ages (young, 35:65; aged, 15:85). The total count matrix was split by random sampling into a 10% held out validation set and a 90% train/test set. “Chrono-variant” genes were identified based on *only* the train/test set to avoid information leakage from the validation set. All genes that showed a *>* 0.1 fold change on a natural log scale and were expressed in at least 3 cells were considered chrono-variant, yielding 667 genes. SVM classification models were trained with *L*_1_ regularization to enforce sparsity. Regularization strength *λ ∼ C^−^*^1^ was optimized by performing a line search using 5-fold cross-validation within the train/test set. The number of non-zero weight coefficients in each trained, *L*_1_-regularized classifier was considered to be the number of genes utilized by that classifier. Validation accuracies were obtained by training a classifier on the entire train/test set with the optimized regularization strength, and performing prediction on the held-out validation set.

### 4.14 RNA Velocity Analysis and Dynamical Simulations

RNA velocity was inferred using velocyto [31] with default parameters. Gene expression levels were first imputed using a k-nearest neighbors approach, as outlined [31]. The magnitude of RNA velocity relative to pseudotime was quantified by binning cells along the pseudotime axis and computing the magnitude of the mean RNA velocity for each bin.

A *k*-nearest neighbors regression model (kNN-R) was trained on PCA embeddings for experimentally measured single cell transcriptomes and their corresponding pseudotime assignments. Using 5-fold cross validation, a range of values for *k* were estimated and *k* = 30 was chosen to optimize the regression *r*^2^while minimizing computational expense. *k*-NNR model fit was estimated at *r*^2^ > 0.96 by 5-fold cross validation, indicating high performance for estimation.

To determine differences between aged and young velocity fields, phase point simulations were performed with numerical methods. A set of initial positions in the 2D PCA embedding for both young and aged cells was sampled from observed cellular positions. For these experiments, the set of initial positions was restricted to cells in the “activated” 18 hour time point to prevent simulations from encountering the low density region between quiescent and activated cells where we have little information for velocity inference. Additionally, we restrict initial positions to observed cells with a PC1 embedding score *< −*3, which corresponds to the more primitive cells in transcriptional Cluster 2.

Phase points were initiated at positions *x*_0_ and evolved for *t* = 5, 000 timesteps. At each timestep, phase point velocity *v_t_* was computed as the mean velocity of the *k* = 100 nearest cells to the phase point in the observed cell embeddings. For simulations in young and aged velocity fields, only young or aged cells were considered at this step, respectively. New phase point positions *x_t_*_+1_ were computed as the sum of the velocity *v_t_* and current phase point position *x_t_*, plus a noise term *η*:

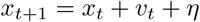

where noise is drawn from a multidimensional normal distribution *η* (0, 𝒩*σ_t_*) with a standard deviation *σ_t_* computed as the standard deviation of the velocity from the *k* = 100 nearest cells to the phase point. When specified, this noise term was set to 0 for some experiments. Code is available in the MuSC Atlas Github.

### 4.15 Change in Pseudotime Analysis

The “change in pseudotime” (ΔPseudotime) was estimated for each cell using the *k*-nearest neighbors regression model. Future transcriptional states *x_t_*_+1_ were inferred by RNA velocity as above, and the pseudotimes for these states were predicted using the kNN-R model. ΔPseudotime is defined as the difference between the inferred future and measured present pseudotime for each cell:

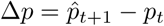

where *p̂_t_*_+1_ is the inferred pseudotime using RNA velocity and the kNN-R model and *p_t_* is the observed pseudotime at the experimental timepoint.

Cells were defined to be undergoing “lineage regression” if they displayed a ΔPseudotime *< σ*, where *σ* is the standard deviation of the ΔPseudotime distribution. Code is available in http://github.com/jacobkimmel/myodyn.

## 5 Authors Contributions

JCK conceptualized the study, performed timelapse imaging and single cell RNA-sequencing experiments, analyzed data, and wrote the paper. AH assisted with single cell RNA-sequencing and flow cytometry experiments. WFM and ASB provided funding, supervised research and edited the manuscript.

## 6 Acknowledgements

The authors would like to thank Kurt Thorn, DeLaine Larsen, and Andrew G. York for helpful discussions.

## 8 Data Availability

Raw and processed single cell RNA-seq data have been submitted to NCBI GEO. Single cell RNA-seq associated code and cell motility tracks are available at http://github.com/jacobkimmel/myodyn. Heteromotility is available at http://github.com/cellgeometry/heteromotility, our segmentation models are available at http://github.com/jacobkimmel/fcdensenet pytorch, and our cell tracking code is available at http://github.com/jacobkimmel/musc tracker.

## 10 Supplemental Information

### 10.1 Supplemental Figures

**Figure S1:**
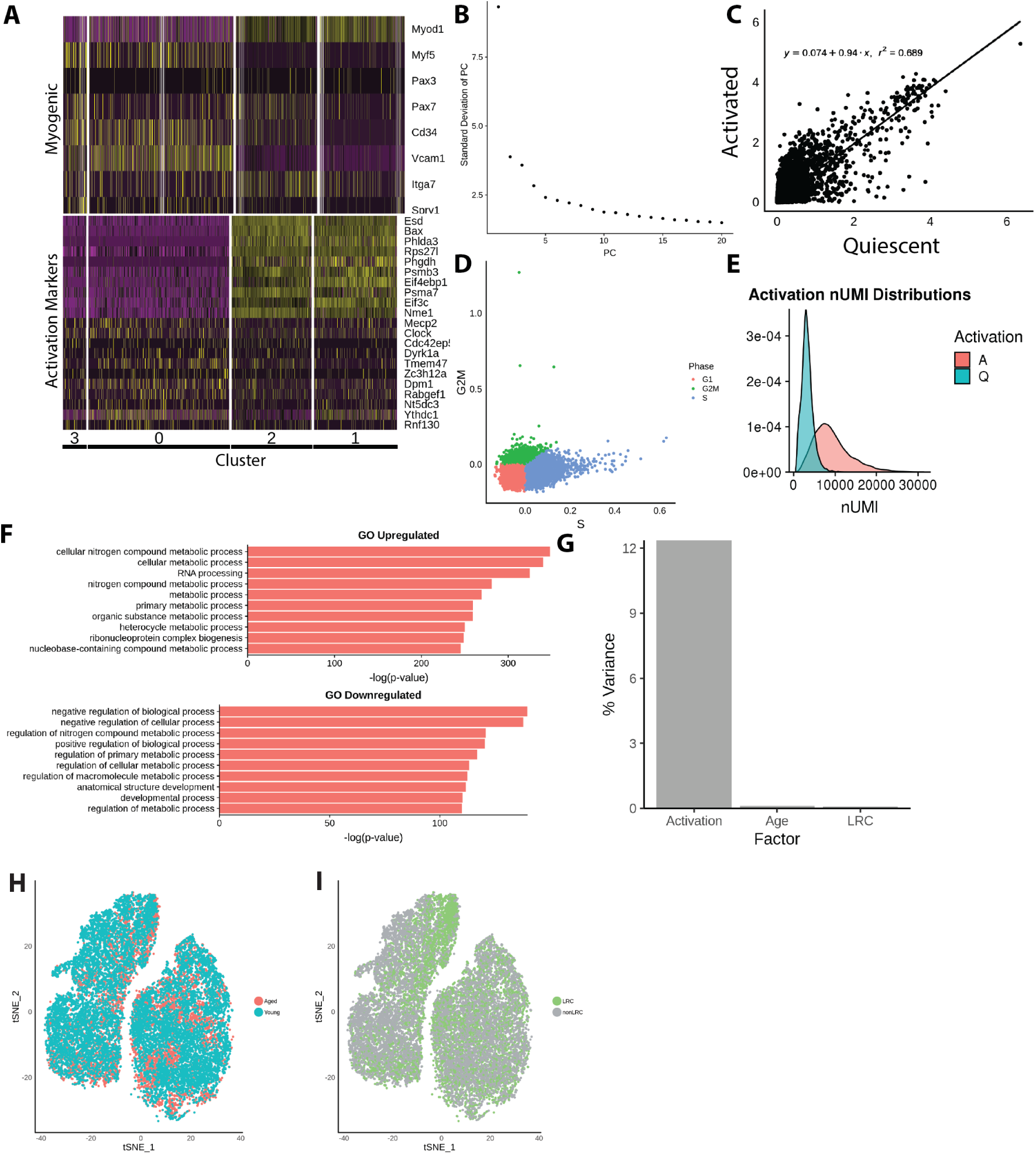
Myogenic activation engages biosynthetic transcriptional programs. **(A)** Heatmap of myogenic genes and activation markers across transcriptional clusters. **(B)** Top 10 positive and negative loadings for the first two principle components. **(C)** Gene-wise correlation for mean expression levels between quiescent and activated MuSCs. **(D)** Cell Cycle scoring demonstrating little difference in cycle state across our population. **(E)** Distributions of total UMI counts per cell for quiescent and activated MuSCs, demonstrating a global increase in mRNA content for activated cells. **(F)** GO enrichment analysis for genes upregulated and downregulated in activation relative to quiescence. **(G)** Proportion of variation in transcriptomes explained by each of the 3 factors in the single cell RNA-seq experiment: Age, proliferative history (LRC status), and Activation state (time in culture, 0 hr or 18 hr). Activation is a much greater source of variation than aging or proliferative history.

**Figure S2:**
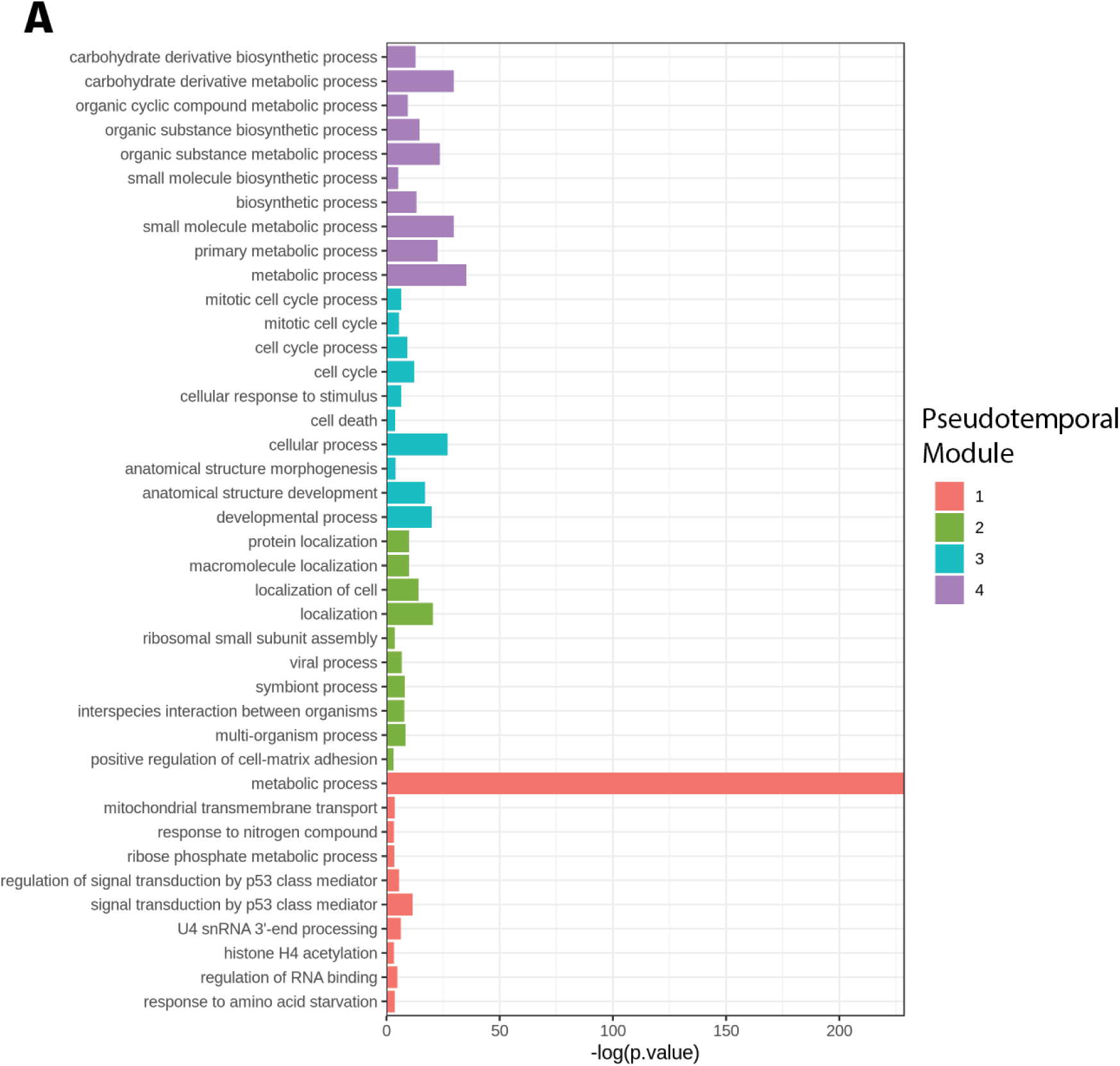
Pseudotemporal analysis of myogenic activation reveals non-monotonic regulation. **(A)** Gene ontology enrichment analysis for Pseudotemporal Modules, suggesting coherent groups of co-regulated genes.

**Figure S3:**
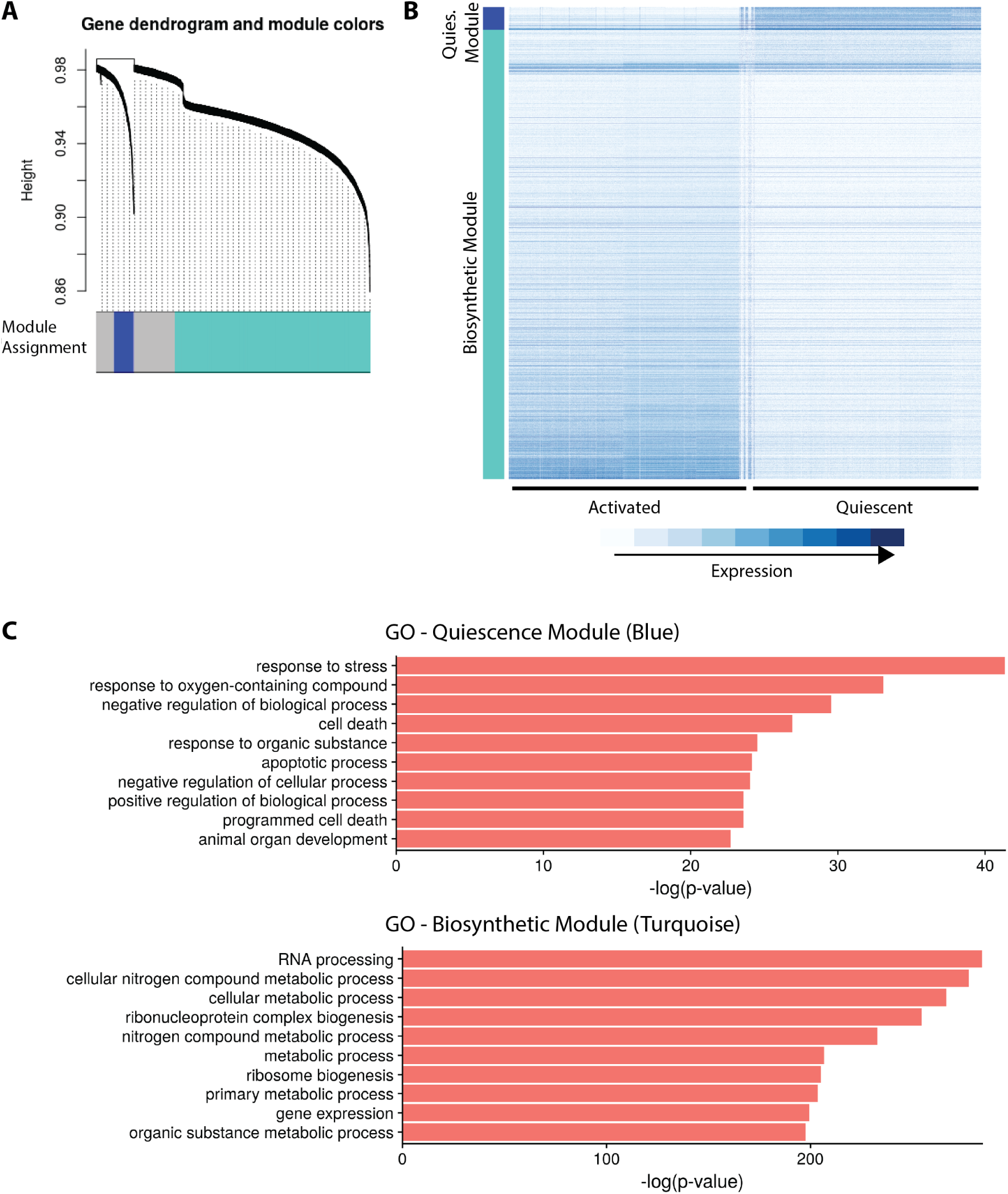
Weighted gene correlation network analysis. **(A)** Module identification in the weighted gene correlation network, with significant modules indicated by colored labels. **(B)** Heatmap of genes in the identified Quiescence and Biosynthetic modules in quiescent and activated cells. **(A)** Gene ontology enrichment analysis for genes in the identified Quiescence and Biosynthetic modules. The Quiescence module is notably enriched for stress response and cell death regulation genes, while the Biosynthetic module is© cenriched for RNA and protein biosynthesis.

**Figure S4:**
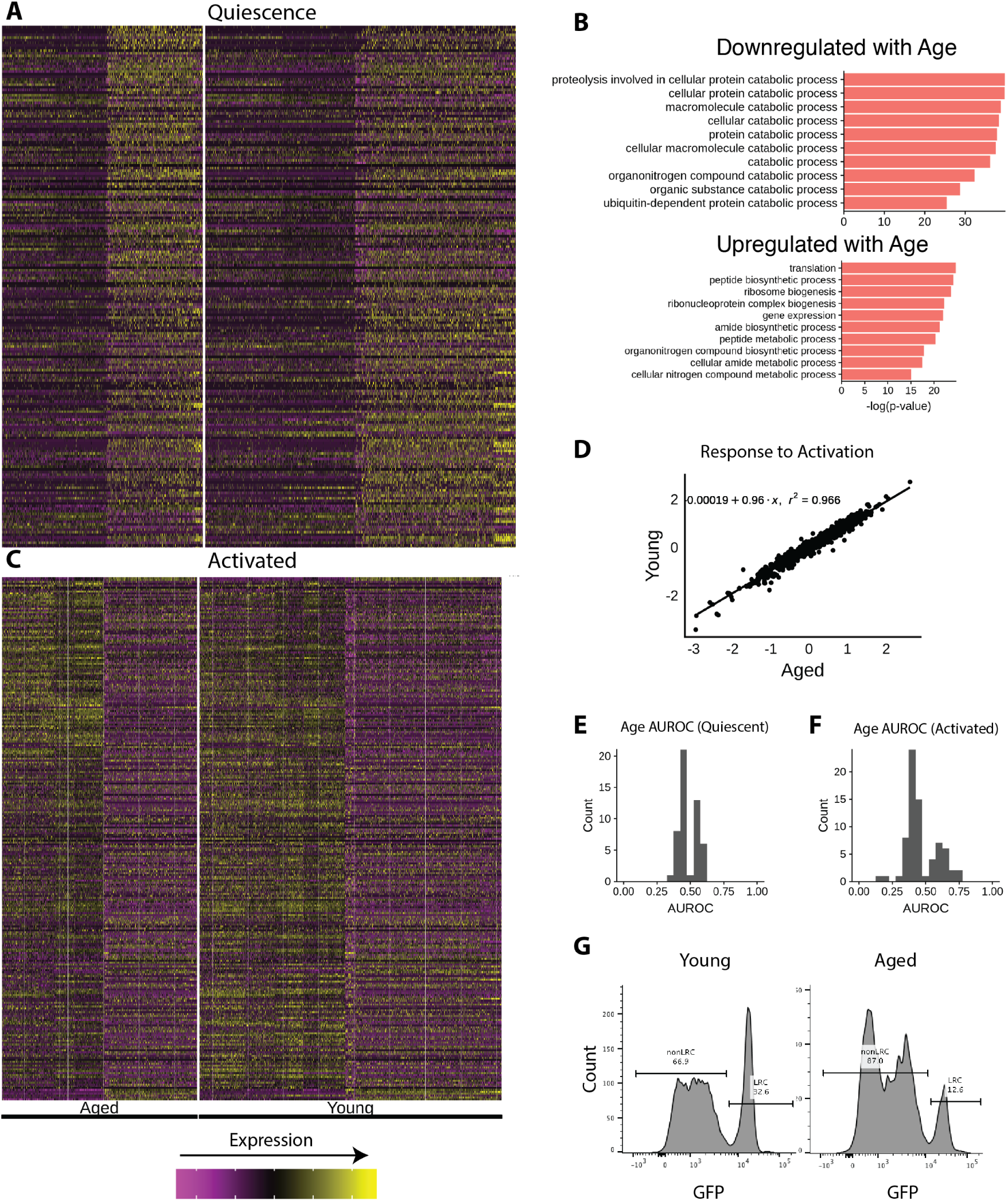
Aged MuSCs display differential activation responses in many genes. **(A)** Heatmaps of differentially expressed genes between aged and young quiescent MuSCs, and **(C)** activated MuSCs. As indicated by the heatmaps, there are no individual markers with high discriminatory ability, and activation increases transcriptional differences between aged and young MuSCs. **(B)** Gene ontology enrichment analysis for differentially expressed genes between aged and young activated MuSCs. Stress responses and cell death processes are upregulated in aged cells. **D** Genewise correlation for transcriptional differences induced by activation (Activated mRNA Count - Quiescent mRNA Count) between aged and young MuSCs. **E** Distances between aged and young MuSC centroids in PC space when LRC:nonLRC ratios are subsampled to be equal (35% LRC, “AYD-Equal”) or subsampled to reflect physiological differences between aged and young LRC ratios (10% LRC in aged, 35% in young, “AYD-Phys”). Distances between randomly selected aged (“Aged-Null”) or young (“Young-Null”) are presented for comparison. **F** Distribution of age classification AUROC scores for individual genes in quiescent and **(G)** activated MuSCs. **(H)** Representative flow cytometry measurements of the proportion of LRC (GFP high) and nonLRC (GFP low) MuSCs.

**Figure S5:**
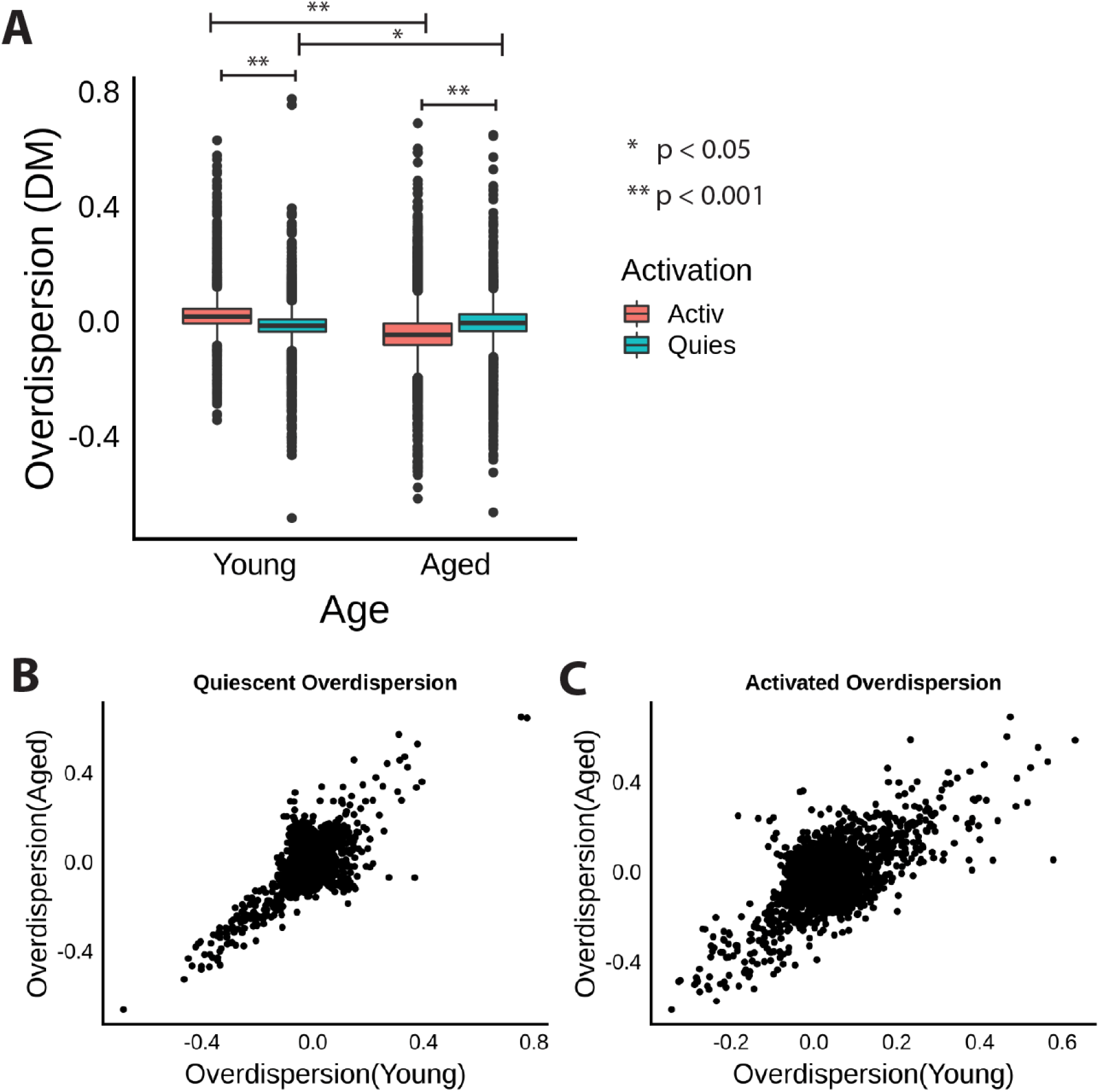
Aging changes gene expression variation in a context dependent manner. **(A)** Overdispersion distributions for young and aged MuSCs in both quiescent and activated conditions. Each underlying point represents the overdispersion estimate for a single gene. Gene expression variance increases with activation in young cells, but decreases with activation in old cells. **(B)** Comparison of overdispersion estimates for each gene between young and aged cells in quiescent and **(C)** activated conditions. in quiescent and activated cells.

**Figure S6:**
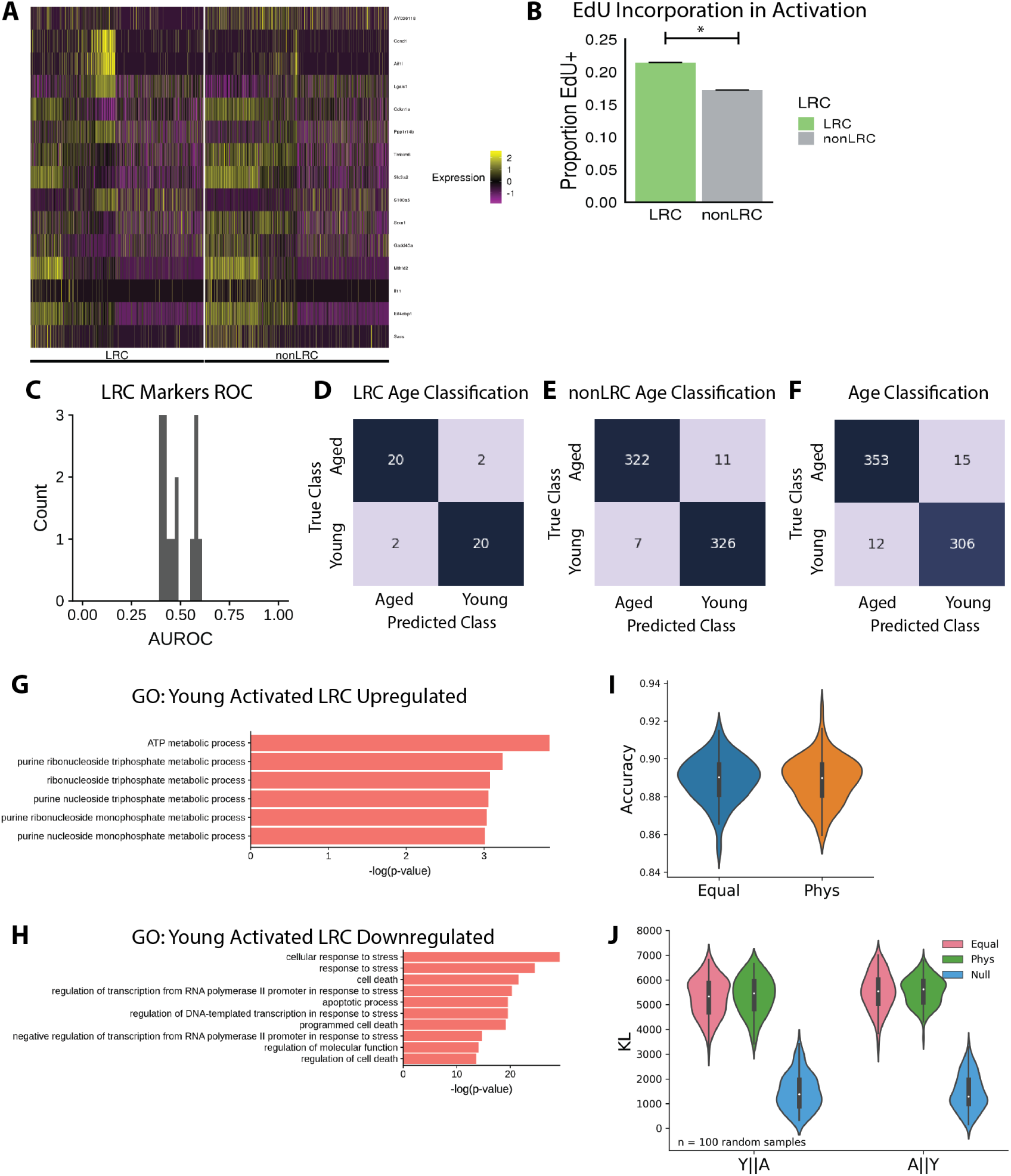
LRCs and nonLRCs are discriminated by small differences in many genes. **(A)** Differentially expressed genes between LRCs and nonLRCs in both quiescent and activated states. **(B)** EdU incorporation by LRCs and nonLRCs during early activation in culture. **(C)** Distribution of LRC classification AUROC scores for individual genes. **(D)** Confusion matrices for classification of age in activated LRCs, **(E)** nonLRCs, and **(F)** the total MuSC pool using our feature selection pipeline and support vector machine classifier. **(G)** GO enrichment analysis in young MuSCs for terms upregulated in LRCs and **(H)** downregulated in LRCs. **(I)** Classification accuracy for a probabalistic neural network classifier trained to discriminate young vs. aged MuSCs with either (1) equal ratios of LRC:nonLRC (35:65) at both ages or (2) physiologically observed ratios (35:65 in young, 15:85 in aged). Classification accuracy is equal between the two, suggesting that a change in LRC ratios is a minor contribution to the “magnitude” of aging. Accuracies presented are the mean of a 5-fold cross-validation split. **(J)** Estimated Kullback-Leibler divergence between young and aged MuSCs in random samples with either (1) equal LRC:nonLRC ratios (35:65) or (2) physiologically observed ratios. The divergence is equal in both contexts in both directions (the KL divergence is asymmetric) suggesting that changes in LRC proportions do not dramatically alter the magnitude of transcriptional change with age.

**Figure S7:**
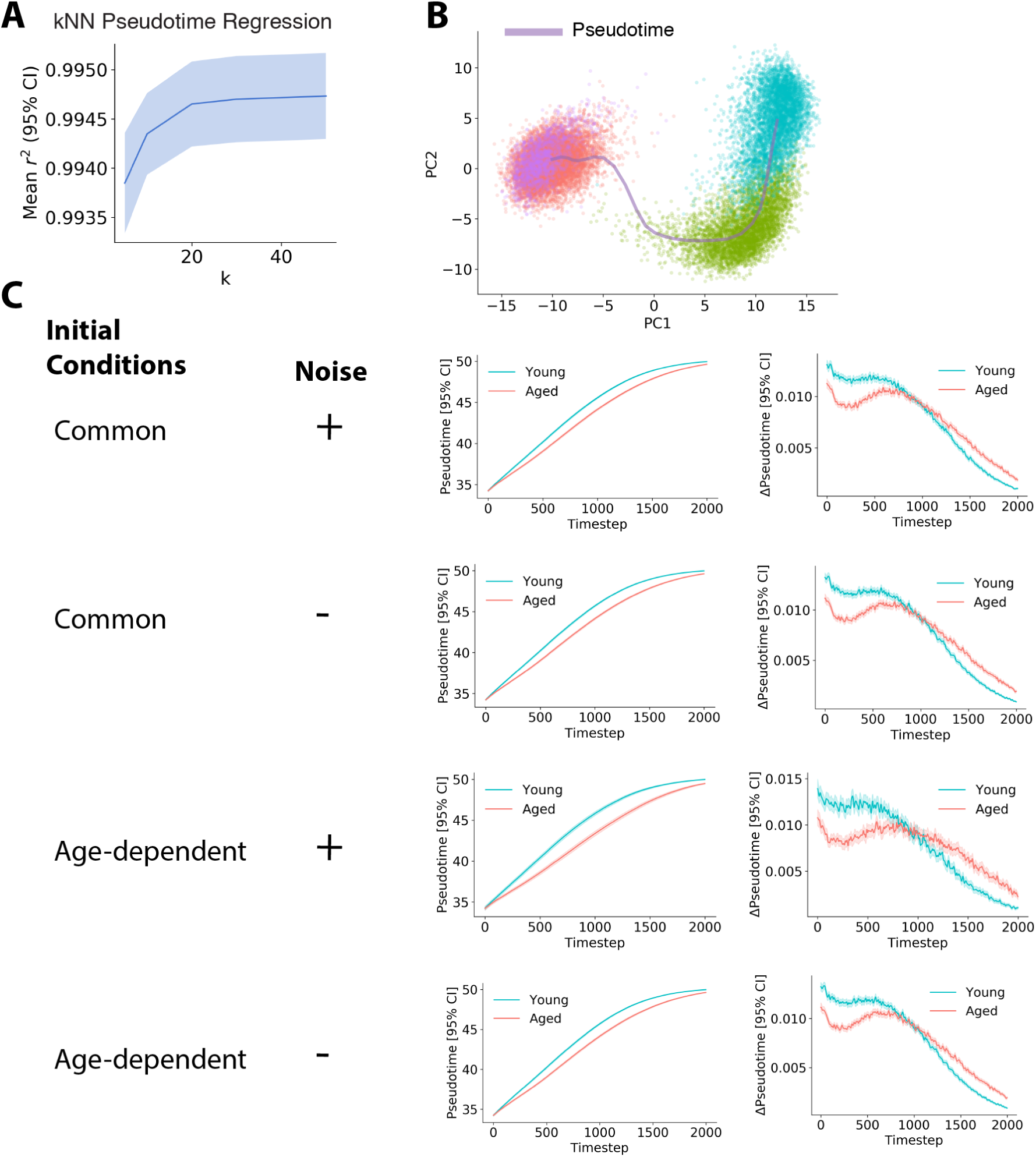
Phase point simulations reveal differences in state dynamics between aged and young MuSCs. **(A)** Prediction accuracy measured using Pearson’s *r*^2^ for a *k*-nearest neighbors regression model mapping transcriptional principal component scores to pseudotime values. Validated with 5-fold cross-validation. Data are presented as the mean and 95% confidence interval. **(B)** Visualization of the pseudotime curve in PCA space, estimated by computing a rolling mean of PC scores across the 1D pseudotime coordinate. Colors represent transcriptional state clusters. **(C)** Progression through pseudotime for young and aged phase points simulated with varying model parameters. Initial conditions were either (1) “common,” sampled from all observed cell positions in transcriptional space, with each initial condition simulated using aged or young velocities, or (2) “age-dependent,” where all intial conditions simulated using young velocities were sampled exclusively from positions observed among young cells, and vice-versa. Simulations either included Gaussian noise *η* scaled by the standard deviation of velocity in the local neighborhood, or no noise.

**Figure S8:**
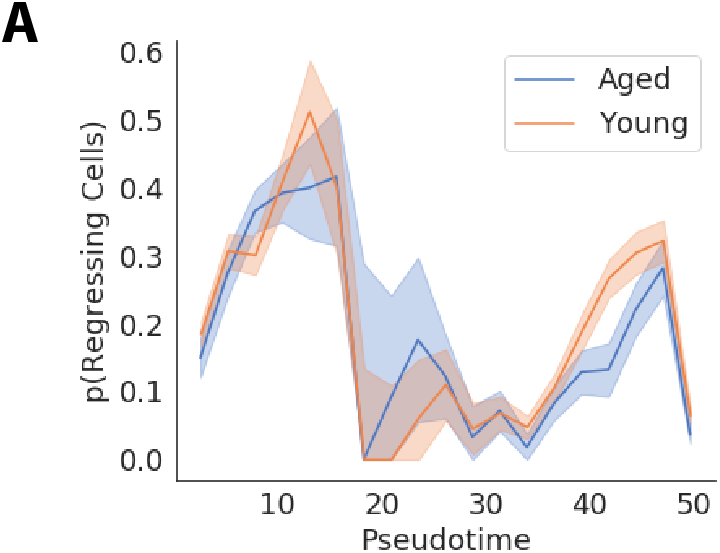
Lineage regression frequency with age. **(A)** The proportion of “regressing” cells across pseudotime in aged and young MuSCs, revealing little age dependence.

### 10.2 Supplemental Tables

**Table S1:** Cell counts by condition.

| Activation | Age | LRC | Cell Count |
| --- | --- | --- | --- |
| A | Aged | LRC | 495 |
| A | Aged | nonLRC | 3362 |
| A | Young | LRC | 3236 |
| A | Young | nonLRC | 3636 |
| Q | Aged | LRC | 1074 |
| Q | Aged | nonLRC | 2697 |
| Q | Young | LRC | 3507 |
| Q | Young | nonLRC | 3548 |

### 10.3 Supplemental Files

**Movie S1** Heatmaps representing the density of phase points over time in transcriptional space during phase analysis simulations in young (left) and aged (right) velocity fields. Transcriptional space is represented using the first two principal components. Darker colors indicate higher cell densities for both color maps.

